# Clathrin-independent endocytosis of Human Papillomaviruses is facilitated by actin nucleation promoting factor WASH

**DOI:** 10.1101/2021.06.18.448076

**Authors:** Pia Brinkert, Lena Krebs, Pilar Samperio Ventayol, Lilo Greune, Carina Bannach, Delia Bucher, Jana Kollasser, Petra Dersch, Steeve Boulant, Theresia E.B. Stradal, Mario Schelhaas

## Abstract

Endocytosis is a fundamental cellular process facilitated by diverse mechanisms. Remarkably, several distinct clathrin-independent endocytic processes were identified and characterized following virus uptake into cells. For some, however, mechanistic execution and biological function remain largely unclear. This includes an endocytic process exploited by human papillomavirus type 16 (HPV16). Using HPV16, we examined how vesicles are formed by combining systematic cellular perturbations with electron and video microscopy. Cargo uptake occurred by uncoated, inward-budding pits. Mechanistically, vesicle scission was facilitated by actin polymerization controlled through the actin nucleation promoting factor WASH. While WASH typically functions in conjunction with the retromer complex on endosomes during retrograde trafficking, endocytic vesicle formation was largely independent of retromer itself and the heterodimeric membrane-bending SNX-BAR retromer adaptor, thereby uncovering a role of WASH in endocytosis in addition to its canonical role in intracellular membrane trafficking.

## Introduction

Cells internalize extracellular ligands, fluids, plasma membrane lipids, and receptors by endocytosis. It is crucial for cellular and organismal homeostasis or responses by for example regulating signal transduction, intercellular communication, and neurotransmission (Schmid and Conner, 2003). Several endocytic mechanisms exist that are distinguished by diverse criteria. Morphologically, cargo is engulfed either by outward membrane protrusions (macropinocytosis, phagocytosis) or by inward budding pits (e.g. clathrin-mediated endocytosis (CME), glycosphingolipid enriched endocytic carriers (GEEC)) (Heuser and Evans, 1980; Kirkham et al., 2005). The cellular machinery executing endocytic vacuole formation further defines the identity of endocytic pathways (Doherty and McMahon, 2009). For instance, macropinocytosis occurs by growth factor-induced, actin-driven protrusions that form large vacuoles upon backfolding with help of the Bin1/Amphiphysin/Rvs (BAR) protein C-terminal binding protein 1 (CtBP1) (Liberali et al., 2008; Lim and Gleeson, 2011). Of the inward budding mechanisms, CME occurs by sequential recruitment of adaptor proteins (AP) such as AP2 and the clathrin coat. Dynamin-2 mediated scission then leads to vacuole formation (McMahon and Boucrot, 2011; Merrifield and Kaksonen, 2014). In contrast, caveolar endocytosis occurs by preassembled coat structures (Pelkmans et al., 2004; Hayer et al., 2010; Stoeber et al., 2016). Another criterium is cargo specificity. For instance, macropinocytosis and CME are responsible for bulk fluid uptake or specific membrane receptor internalization, respectively. In addition to the well-established pathways, a number of less well understood endocytic processes have emerged. Only little information is available on which cellular processes they regulate, or on how vesicles are formed, as they are defined by their independence of classical endocytic regulators such as clathrin, caveolin, dynamin-2, or cholesterol (Doherty and McMahon, 2009). Importantly, several of these mechanisms have been identified by following the internalization of viruses such as lymphocytic choriomeningitis virus (LCMV), influenza A virus (IAV), and human papillomavirus type 16 (HPV16) (Rust et al., 2004; Quirin et al., 2008; Mercer et al., 2010; Schelhaas et al., 2012).

As intracellular parasites, viruses depend on host cells for their life-cycle. This is particularly important during the initial phase of infection termed entry, during which viruses deliver their genome to the site of replication within cells. As virus particles lack locomotive abilities, they strictly rely on cellular transport mechanisms to cross the plasma membrane, the cytosol, or the nuclear envelope. Following viral particles *en route* to the site of replication thus allows to uncover essential cellular mechanisms (Marsh and Helenius, 2006). For instance, early studies on Semliki Forest virus (SFV) coined the field of endosomal trafficking by contributing to the identification of endosomes as pre-lysosomal structures (Helenius et al., 1983). Moreover, research on simian virus 40 (SV40) had large impact on the understanding of caveolar endocytosis (Anderson et al., 1996; Pelkmans et al., 2001; Thorley et al., 2010). More recently, our studies on HPV16 highlighted the existence of a distinct endocytic mechanism independent of most known endocytic regulators (Schelhaas *et al*., 2012).

HPV16 is a small non-enveloped DNA virus and the leading cause of cervical cancer (de Sanjose et al., 2010; Mirabello et al., 2017). It initially infects basal keratinocytes of squamous mucosal epithelia, whereas completion of its life cycle requires suprabasal keratinocyte differentiation (Doorbar, 2005). Uptake of HPV16 into cells is slow and asynchronous, with quick individual virus internalization events occurring over a period of many hours after binding to cells (Schelhaas *et al*., 2012; Becker et al., 2018). Initial binding to heparan sulfate proteoglycans (HSPGs) induces crucial structural changes in the virus capsid that allow transfer to an internalization receptor complex to induce endocytosis (Giroglou et al., 2001; Richards et al., 2006; Selinka et al., 2007; Cerqueira et al., 2013; Cerqueira et al., 2015; Becker *et al*., 2018). While not understood in detail, the existing evidence supports a model in which the receptor complex constitutes specialized tetraspanin-enriched microdomains (TEMs) that resemble hemidesmosomes (HDs) consisting of the tetraspanin cluster of differentiation (CD) 151 (Scheffer et al., 2013; Fast et al., 2018), integrin α6 (Evander et al., 1997; Yoon et al., 2001; Scheffer *et al*., 2013) and epidermal growth factor receptor (EGFR) (Schelhaas *et al*., 2012; Surviladze et al., 2012; Bannach et al., 2020). After endocytosis, HPV16 traffics through the endosomal system (Smith et al., 2008; Spoden et al., 2008; Bienkowska-Haba et al., 2012). From endosomes, the virus is routed to the trans-Golgi network by retrograde transport (Day et al., 2013; Lipovsky et al., 2013) and reaches its site of replication, the nucleus, after nuclear envelope breakdown during mitosis (Pyeon et al., 2009; Aydin et al., 2014).

As indicated, HPV16 endocytosis is independent of a long list of prominent endocytic regulators including clathrin, caveolin, dynamin, and cholesterol (Schelhaas et al., 2012). This endocytic mechanism depends on sodium/proton exchange, actin polymerization, signaling via EGFR, phosphatidylinositol-4,5-bisphosphate 3-kinase (PI3K), p21-activated kinase 1 (PAK1), protein kinase C (PKC), and Abelson tyrosine-protein kinase 2 (Abl2) (Schelhaas *et al*., 2012; Bannach *et al*., 2020). It remains largely elusive how these factors contribute to endocytic vesicle formation. We know, however, that EGFR and Abl2 regulate endocytosis induction and pit maturation, respectively (Bannach *et al*., 2020). As such, the requirements for this endocytic pathway are somewhat reminiscent of macropinocytosis, during which sodium/proton exchange regulates large actin-driven membrane protrusions to engulf bulk extracellular material (Doherty and McMahon, 2009; Mercer *et al*., 2010). However, endocytosis results in small vesicles of about 60-100 nm diameter that are generated independently of cholesterol and the classical Rho-like GTPases cell division cycle 42 (Cdc42), rat sarcoma (Ras)-related C3 botulinum toxin substrate 1 (Rac1), and Ras homolog family member 1 (RhoA) typically associated with macropinocytosis (Schelhaas *et al*., 2012).

Here, addressed the mechanistic execution of vesicle formation using HPV16 as cargo. We showed that cargo uptake occurred via uncoated vesicles severed from the plasma membrane by actin-related proteins (Arp) 2/3 complex-dependent branched actin polymerization. Interestingly, Arp2/3 complex activation occurred by the nucleation promoting factor (NPF) Wiskott-Aldrich syndrome protein (WASP) and suppressor of cAMP receptor (Scar) homologue (WASH), a well-known regulator of endosomal cargo sorting, but not NPFs typically found at the plasma membrane. Distinct from its role at endosomes, WASH was not recruited to endocytic sites by the endosomal retromer complex or the membrane bending sorting nexin (SNX) hetero-dimer. Moreover, WASH function was independent of ubiquitylation at the established site K220, but required phosphorylation at Y141. Thus, our findings demonstrated a direct involvement of WASH in endocytosis distinct from its canonical role at endosomes.

## Results

### Endocytic vesicle formation occurred via uncoated, inward budding vesicles

Since the mechanism of how endocytic vesicles are formed is mostly negatively and thus ill-defined, we initially set out to elucidate its morphological itinerary. HPV16 was employed as trackable cargo in correlative light and electron microscopy (CLEM). To visualize different stages of endocytic vacuole formation and potential coats lining the forming vesicle as seen for clathrin-coated pits or caveolae, fluorescent virus particles were correlated with structures on the cytosolic leaflet of plasma membrane sheets analyzed by transmission EM of metal replicas (Heuser and Evans, 1980; Sochacki et al., 2014; Bucher et al., 2018). As inherent quality control, different stages of clathrin-coated pit formation were readily observable at sites distinct from virus localizations (Figure 1A). For HPV16, about 20% of viruses were associated with no obvious structure, likely representing virus binding to HSPGs prior to engagement of specialized TEMs and endocytosis induction. A considerable proportion (31%) was correlated with dense actin network patches (Figure 1B). These may constitute anchoring structures for TEMs, either prior to or during induction of endocytosis (Schelhaas et al., 2008; Ménager and Littman, 2016). Importantly, inward-budding structures at virus sites were readily observed and classified into stages of vesicle formation in analogy to CME (Figures 1A and 1B). Designated as early stage, 14% of virions were associated with roughened plasma membrane patches representing initial curvature formation (Figures 1A and 1B). Small invaginations (100-150 nm in diameter) were assigned as intermediate stage, and represented pit maturation (13%). Fully rounded invaginations of 80-100 nm (22%) were classified as late-stage endocytic pits ready for scission (Figures 1A and 1B). Notably, all invaginations were devoid of a discernable coat or other regular structures (Figure 1A). Thus, cargo uptake in the unique endocytic pathway occurred by a stepwise, inward budding process, in which endocytic vacuoles were formed without observable contribution of a coat structure. One can infer from the stepwise formation of vesicles that different sets of proteins must be recruited and function in a sequential manner to facilitate membrane invagination, pit dilation, constriction and scission in analogy to CME.

**Figure 1.**
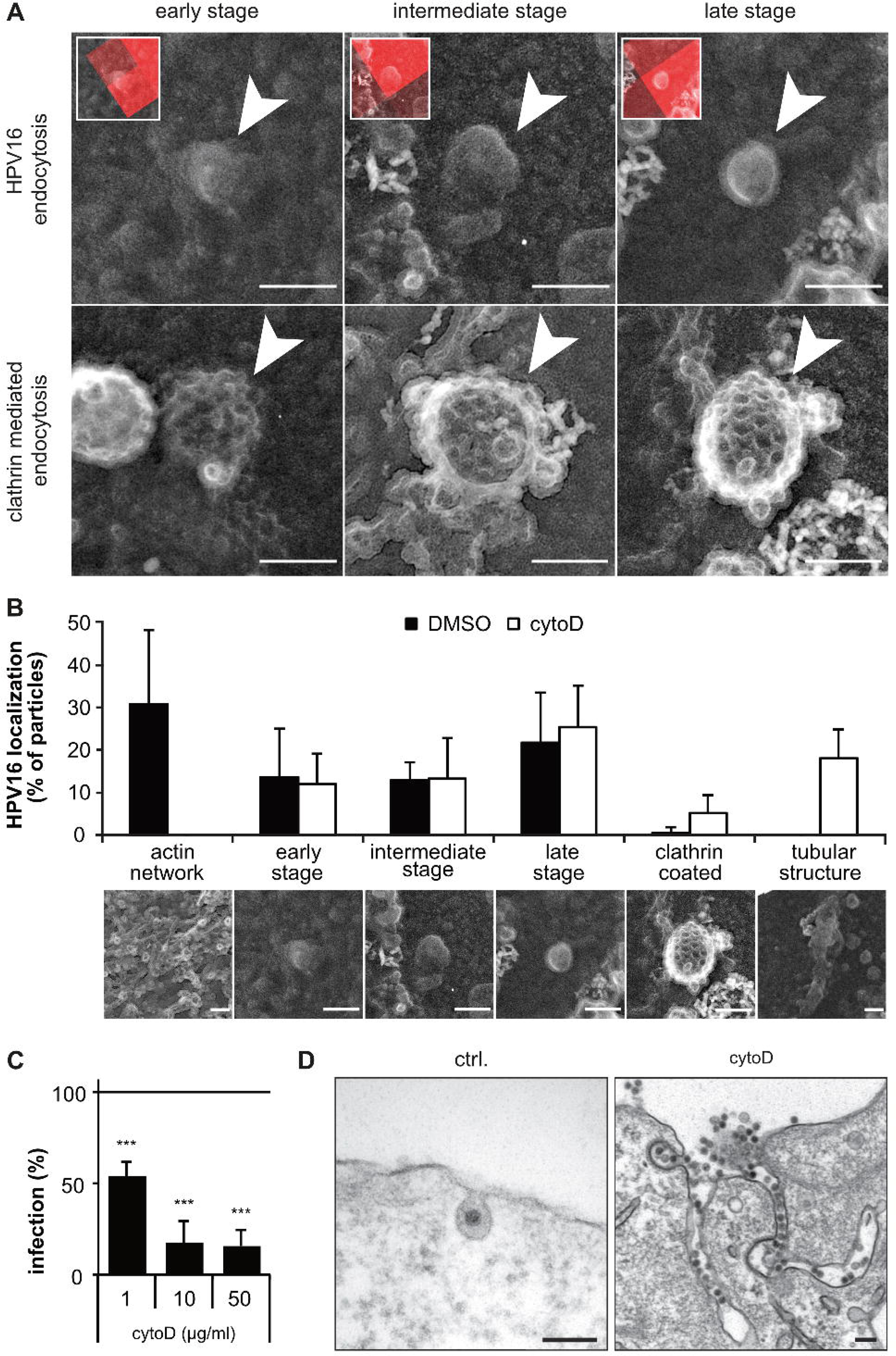
HPV16 endocytosis is an uncoated, inward budding mechanism. (A) HaCaT cells were seeded on HPV16-AF568 bound to ECM, treated with 10 µg/ml cytoD or DMSO and unroofed. Fluorescent virus particles imaged by spinning disk microscopy and depicted in the small insets were correlated with structures identified in transmission EM images of platinum/carbon replicas of unroofed membranes. Arrowheads indicate endocytic pits. (B) Depicted is the percentage of particles associated with the indicated structure ± variation between the membranes (DMSO: 134 viruses/ 7 membranes; cytoD: 101 viruses/5 membranes). Of note, about 20-25% of particles were not associated with any prominent structure. (C) HeLa ATCC cells were infected with HPV16-GFP in presence of cytoD. Infection was scored by flow cytometry and normalized to DMSO treated controls. The mean of three experiments ± SD is shown. (D) HeLa ATCC cells treated with 10 µg/ml CytoD or left untreated (ctrl.) were infected with HPV16-GFP and processed for ultra-thin section transmission EM at 6 h p.i.. All scale bars are 100 nm.

In the studied mechanism, most typical endocytic regulators including dynamin are dispensable for vesicle formation. However, actin polymerization facilitates cargo uptake (Schelhaas *et al*., 2012). Accordingly, perturbation of actin assembly by cytochalasin D (cytoD) reduced HPV16 infection (Figure 1C). To analyze how defective actin assembly impacts the steps of endocytic pit formation, we used CLEM of cytoD-treated cells. As expected, the actin network population was no longer detectable (Figure 1B). However, all stages of membrane invaginations were present to a similar extend in cytoD- and untreated samples (Figure 1B) indicating that endocytosis induction and pit formation were independent of actin polymerization. Notably, virus-correlated tubular structures devoid of an apparent coat newly appeared in cytoD-treated cells (Figure 1B). In fact, these tubular membrane invaginations were filled with virus particles (Figure 1D). Taken together, the formation of endocytic structures and the failure to separate cargo-filled tubules from the plasma membrane strongly imply that endocytic vesicle scission but not induction and membrane invagination were facilitated by polymerizing actin filaments.

### Actin polymerization coincided with cargo internalization

Consistent with actin polymerization aiding vesicle scission, the presence of actin was confirmed at the neck of constricted endocytic pits in immunogold labelling EM (Figure 2A). To strengthen this notion and to gain insights into how actin may facilitate vesicle scission, we employed live cell TIRF microscopy (TIRF-M) to selectively illuminate the basal plasma membrane. Denoted by a rapid loss of virus signal, our previous work already demonstrated that vesicles are formed within about 2 minutes, once HPV16 has been committed to uptake (Schelhaas *et al*., 2012). Here, cargo uptake into the cell interior correlated with an increase of filamentous actin, indicating actin polymerization at the time of scission (Figures 2B and 2C, Suppl. Movie 1). Detailed analysis of intensity profiles revealed that the loss of virus signal occurred for about 4 s (Figures 2C and 2D, Suppl. Movie 1). While the onset of actin polymerization was somewhat variable, the increase in actin signals was initiated always prior to cargo internalization (about 10 s on average), peaked just before cargo uptake, and decreased thereafter (Figures 2C and 2D, Suppl. Movie 1). The dynamics of actin polymerization resembled dynamin recruitment during scission in CME, starting about 20 s prior to loss of the clathrin signal from the plasma membrane (Figures S1A-S1C, Suppl. Movie 2) (Merrifield et al., 2002). Since dynamin is dispensable for cargo uptake in this pathway (Spoden *et al*., 2008; Schelhaas *et al*., 2012), actin likely replaced dynamin functionally as a scission factor. Moreover, the local, transient increase in filamentous actin indicated that polymerized actin did not merely serve as an anchor for other scission factors, but was specifically induced for and actively contributed to endocytic vesicle scission as for other endocytic mechanisms (Merrifield *et al*., 2002; Pelkmans et al., 2002; Mooren et al., 2012).

**Figure 2.**
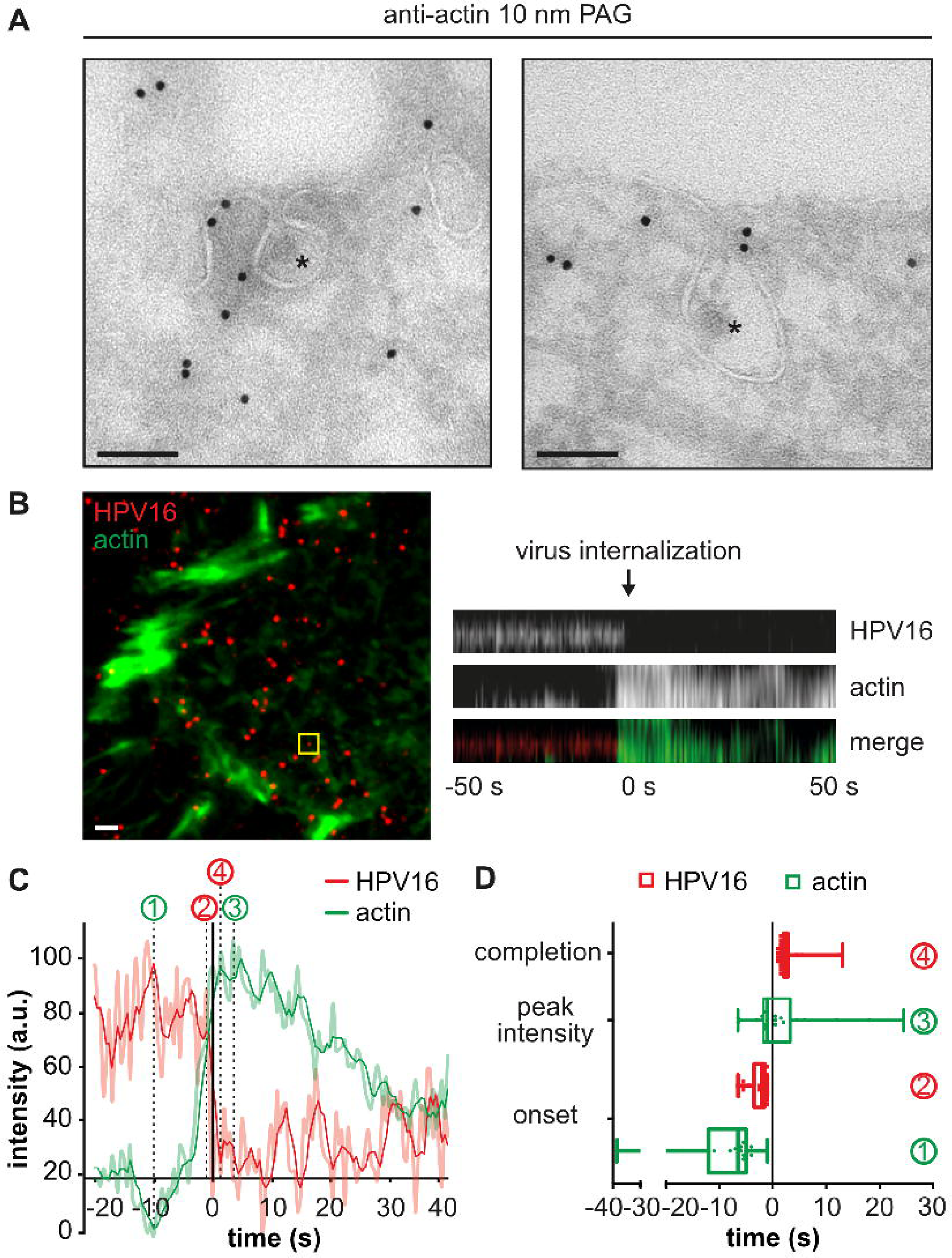
Actin polymerization facilitates cargo internalization. (A) HeLa ATCC cells were infected with HPV16-GFP, subjected to immunogold labeling of actin on ultra-thin cryosections, and analyzed by transmission EM. Asterisks indicate HPV16 in endocytic pits. Scale bars are 100 nm. (B) HeLa ATCC cells were transfected with lifeact-EGFP, infected with HPV16-AF594 and imaged by live cell TIRF-M at 1 h p.i.. Movies were acquired with 0.5 Hz frame rate for 5 min. HPV16 entry events were identified manually after background subtraction and filtering. The yellow box indicates the virus entry event shown as a kymograph (right) and intensity profile (C). Scale bar is 2 µm. (C) Plotted are the intensity profiles of HPV16 and lifeact (light red/green) as well as moving averages (intense red/green). Values are depicted relative to the half time of virus loss (t = 0). The time points of the onset of actin polymerization (1) and of virus signal loss (2) from the cell surface, as well as of actin peak intensity (3) and of completion of virus uptake (4) were determined manually. (D) Time points were determined for 21 virus entry events as indicated in (C), averaged and are depicted as box plots.

### Branched actin polymerization regulated cargo uptake

To understand the mechanism of actin polymerization for endocytic vesicle scission, we investigated the involvement of branched versus unbranched actin filaments. Formin-mediated unbranched actin polymerization was perturbed by SMIFH2, a small molecule inhibitor binding the formin homology 2 domain in formins (Rizvi et al., 2009). SMIFH2 treatment dose dependently interfered with actin-dependent CME, as evidenced by a decrease in vesicular stomatitis virus (VSV) infection (Figure 3A) (Cureton et al., 2009). In contrast, SMIFH2 did not affect HPV16 infection (Figure 3A). Furthermore, HPV16 infection was largely unaffected by siRNA-mediated depletion of individual formins (Figure S2A). However, sequestering the Arp2/3 complex as key regulator of branched actin polymerization by overexpression of the Arp2/3 binding domain of WASP and WASP-family verprolin-homologous protein (WAVE) (Scar-WA) (Machesky and Insall, 1998) strongly reduced HPV16 infection compared to cells expressing a control lacking the Arp2/3 binding domain (Scar-W) (Figure 3B). In line with a requirement of branched actin polymerization, RNA interference (RNAi) with Arp3 expression reduced infection by about 80% (Figure 3C) compared to cells transfected with a non-targeting control (ctrl.). Similarly, vaccinia virus (VV) infection by macropinocytosis (Mercer and Helenius, 2008; Mazzon and Mercer, 2014) was reduced upon Arp3 depletion, whereas Semliki Forest virus (SFV) uptake by actin-independent CME (Marsh and Helenius, 1980; Marsh et al., 1984; DeTulleo and Kirchhausen, 1998) was even increased (Figure 3C). Taken together, active branched but not unbranched actin polymerization was crucial.

**Figure 3.**
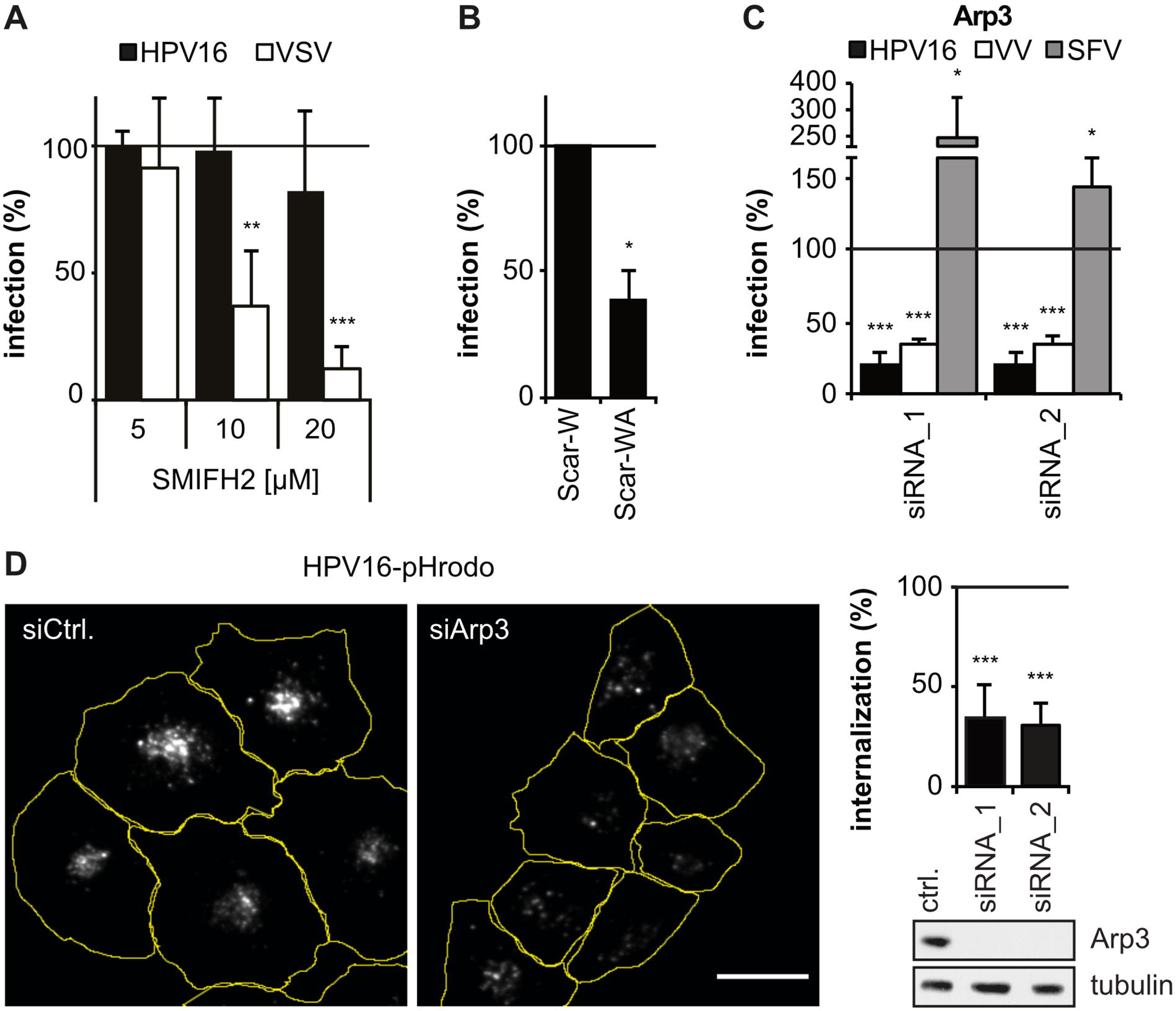
Branched actin polymerization mediates endocytosis. (A) HeLa ATCC cells were infected with HPV16-GFP or VSV-GFP in presence of the formin inhibitor SMIFH2. Infection was scored by flow cytometry, normalized to solvent treated controls and is depicted as mean ± SD (HPV16: n = 3, VSV: n = 4). (B) HeLa ATCC cells transfected with Scar-W-GFP or Scar-WA-GFP were infected with HPV16-RFP. Infection of transfected cells was analyzed by microscopy, normalized to Scar-W-GFP control cells and is depicted as the mean of three experiments ± SD. (C) HeLa Kyoto cells were depleted of Arp3 and infected with HPV16-GFP, VV-GFP or SFV. Infection was scored by automated microscopy or flow cytometry, normalized to cells transfected with a non-targeting control siRNA (ctrl.) and is depicted as mean ± SD (HPV16/SFV: n = 3, VV: n = 4). (D) Arp3 depletion was followed by infection with HPV16-pHrodo and live cell spinning disk microscopy at 6 h p.i.. Shown are average intensity projections of the HPV16-pHrodo signal with cell outlines (yellow), scale bar is 25 µm. Virus signal intensities per cell were measured using a CellProfiler pipeline and normalized to ctrl. (n = 4, mean ± SD). Knock down levels were analyzed by Western blotting against Arp3.

To directly verify the impact on cargo uptake, we employed a *bona fide* endocytosis assay. For this, HPV16 covalently labelled with the pH-sensitive dye pHrodo was used, which gives rise to fluorescence only upon delivery to acidic endosomal organelles (Figure 3D) (Samperio Ventayol and Schelhaas, 2015; Becker *et al*., 2018). Depletion of Arp3 reduced cargo internalization by about 70% (Figure 3D). Thus, branched actin polymerization was essential for endocytosis.

### WASH was the major NPF in HPV16 infection

Since the Arp2/3 complex is inherently inactive, how is it activated for virus uptake? Several NPFs with distinct cellular functions and localizations regulate Arp2/3 complex activity (Stradal and Scita, 2006; Rottner et al., 2010). To identify which NPF acted here, we systematically employed RNAi against all known NPFs in combination with HPV16 infection. Depletion of the most prominent actin regulators in endocytic processes, neuronal WASP (N-WASP) and WAVE isoforms 1 or 2, (Qualmann and Kelly, 2000; Suetsugu et al., 2003; Chadda et al., 2007), did not alter HPV16 infection (Figures S3A-C, and F). Since N-WASP, WAVE1 and 2 were dispensable, a potential involvement of NPFs typically not found at the plasma membrane was assessed next. Neither depletion of junction mediating and regulatory protein, p53 cofactor (JMY), a regulator of DNA damage response and cell migration (Shikama et al., 1999; Zuchero et al., 2009), nor depletion of WASP homolog associated with actin, Golgi membranes and microtubules (WHAMM), which is involved in the secretory pathway (Campellone et al., 2008), impaired HPV16 infection (Figures S3D-F). However, depletion of WASH, a well-known actin regulator of endosomal cargo sorting (Linardopoulou et al., 2007; Derivery et al., 2009; Duleh and Welch, 2010), strongly reduced HPV16 infection, whereas VV uptake by macropinocytosis was only mildly affected (Figures 4A and S3F). In conclusion, the requirement of WASH as the NPF for HPV16 infection suggested that WASH, rather than NPFs typically found at the plasma membrane, acted in HPV16 uptake.

**Figure 4.**
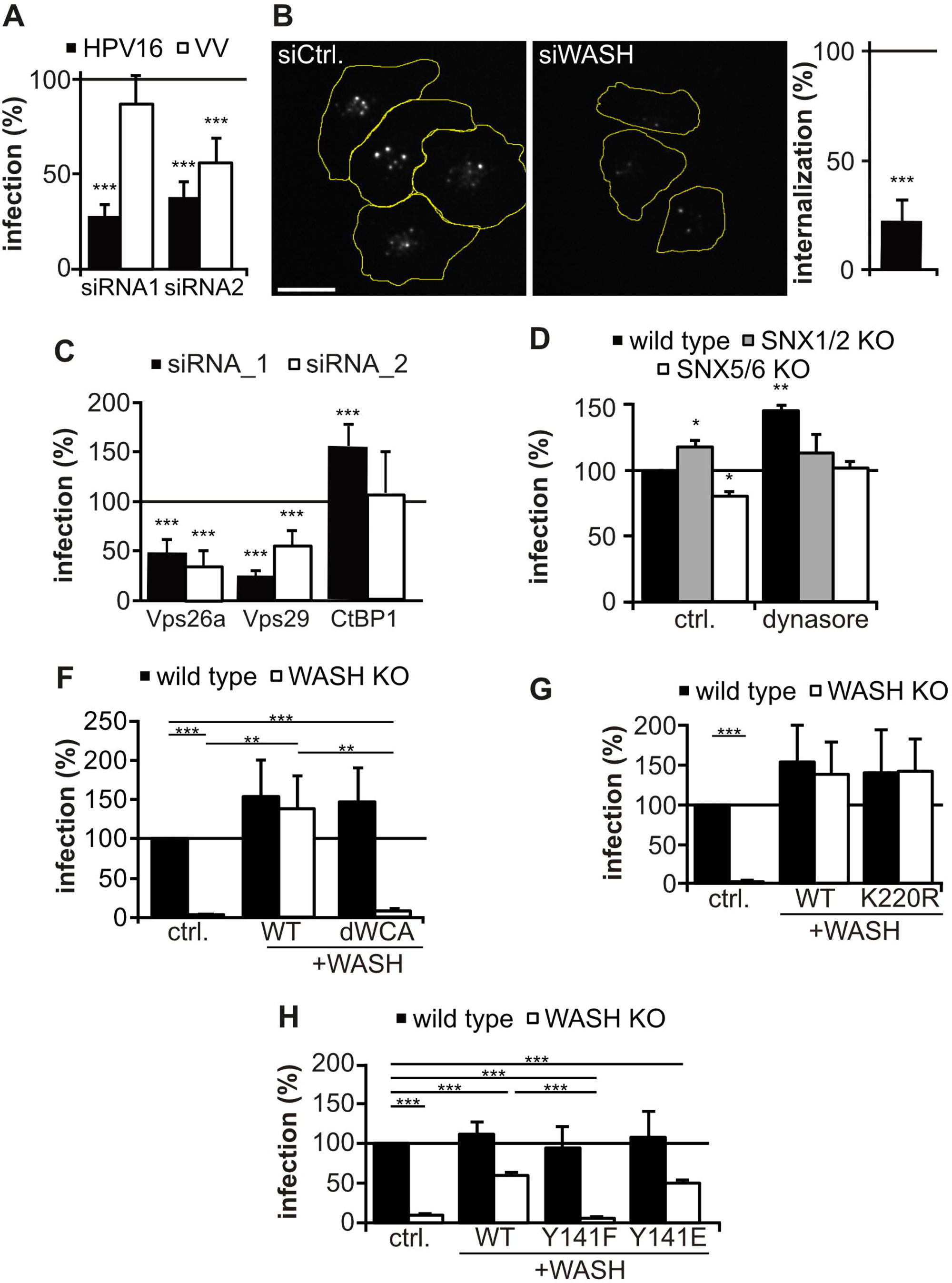
WASH is controlled by Y141 for HPV16 uptake. (A) HeLa Kyoto cells were depleted of WASH and infected with HPV16-GFP or VV-GFP. Infection was determined by automated microscopy and flow cytometry and normalized to cells transfected with a control (ctrl.) siRNA (n = 4, mean ± SD.) (B) After depletion of WASH, cells were infected with HPV16-pHrodo and imaged live by spinning disk microscopy at 6 h p.i.. Shown are average intensity projections of the HPV16-pHrodo signal with cell outlines (yellow), scale bar is 25 µm. The intensity of virus signal per cell was normalized to ctrl. and is depicted as the mean of three experiments ± SD. (C) HeLa Kyoto cells were depleted of Vps26a, Vps29, or CtBP1 and infected with HPV16-GFP. Infection was scored by automated microscopy and normalized to ctrl. (n = 3, mean ± SD). (D) HeLa wild type, SNX1/2 KO or SNX5/6 KO cells were infected with HPV16-GFP. Infection was scored by flow cytometry and is depicted as the mean ± SD (n=3). (E-G) EGFP, WASH-YFP or the WASH mutants dWCA (F), K220R (G) or Y141F/E (H) were expressed in NIH-3T3 wild type and WASH KO cells. Infection with HPV16-RFP was scored by flow cytometry and normalized to wild type cells expressing EGFP (ctrl.). Values are depicted as the mean ± SD (n = 3). Note that (E) and (F) are from the same experiment (same ctrl.) and that WT WASH did not fully restore HPV16 infection in (G) due to shorter expression times compared to (E, F).

### WASH function during virus infection was distinct from its function at endosomes

To our knowledge, no evidence exists that WASH facilitates endocytic vesicle formation. However, since it was the only NPF important for HPV16 infection, we hypothesized a role of WASH in regulating actin polymerization for endocytosis. Consistently, silencing of WASH resulted in a strong decrease of cargo uptake comparable to RNAi of Arp2/3 (Figure 4B, compare Fig 3D). In corroboration, HPV16 infection was completely abrogated in CRISPR/Cas9-derived mouse fibroblast WASH knock-out (KO) cells (NIH-3T3, Figures 4G, H, and S4B), and strongly reduced in human osteosarcoma WASH KO cells (U2OS) (Figures S4A, S4B). This not only confirmed the importance of WASH for cargo uptake, but also underlined the functional existence of WASH-dependent endocytosis in cells derived from different tissues and species.

On endosomes, WASH exerts its function in retrograde transport in conjunction with the retromer complex (Gomez and Billadeau, 2009; Harbour et al., 2012; Jia et al., 2012; Cullen and Steinberg, 2018; Wang et al., 2018; Tu and Seaman, 2021). As previously observed, siRNA-mediated depletion of the retromer proteins vacuolar protein sorting (Vps) 26 and Vps29 interfered with HPV16 infection (Figure 4C) (Lipovsky *et al*., 2013; Popa et al., 2015; Zhang et al., 2018). However, since Vps29 is essential for HPV16 retrograde transport to the Golgi but expendable for virus uptake (Lipovsky *et al*., 2013; Popa *et al*., 2015; Xie et al., 2020), retromer itself is unlikely to facilitate endocytosis. Interestingly, another endosomal protein termed receptor-mediated endocytosis 8 (RME-8) coordinates WASH activity with the membrane bending retromer adaptor SNX-BAR dimer (Freeman et al., 2014). Moreover, recent evidence suggests that the SNX-BAR adaptor consisting of SNX1/2 and SNX5/6 can act independently of the retromer on endosomes (Kvainickas et al., 2017; Simonetti et al., 2017; Simonetti et al., 2019; Yong et al., 2020). Hence, we wondered whether the SNX-BAR dimer and RME-8 play an additional role during endocytosis. As control and to differentiate the mechanism studied here from macropinocytosis, depletion of CtBP1, a prominent BAR domain containing regulator of macropinocytosis (Liberali *et al*., 2008; Valente et al., 2013), did not impair HPV16 infection (Figure 4C). RNAi of RME-8 showed that it was dispensable for HPV16 infection (Figure S5B), To assess the role of the SNX-BAR dimer, we employed CRISPR/Cas knockout cells of SNX1/SNX2 and SNX5/6 (Simonetti *et al*., 2017). Since SNX1/SNX2 and SNX5/6 knockout cells were unimpaired for HPV16 infection (Figure 4D), they were also dispensable. These findings implied that WASH exerted its function during endocytosis in a way distinct from how it typically acts on endosomes.

### WASH acted during endocytosis largely independently from the its regulatory complex at endosomes

Since our findings revealed a distinct WASH function for endocytosis compared to endosomal sorting, we next wondered whether WASH acted in concert with members of the WASH complex (SHRC), namely family with sequence similarity 21 (FAM21), strumpellin, strumpellin- and WASH interacting protein (SWIP), and coiled-coil domain containing 53 (CCDC53). RNAi of the individual members of the complex revealed that while SWIP, strumpellin, and CCDC53 were not needed, FAM21 partially contributed to HPV16 infection (Figure S5F). Hence, WASH acted during HPV16 infection not through the SHRC distinguishing it from its function on endosomes.

### WASH facilitated endocytosis through regulating actin polymerization

This raised the question of whether the function of WASH during HPV16 uptake was facilitated by its function as actin nucleation promoting factor. Ectopic expression of wild type WASH rescued HPV16 infection in WASH knockout cells (Figure 4D-G), whereas expression of WASH lacking the WASP-homology 2, central and acidic (WCA) domain crucial for Arp2/3 complex activation did not (Figure 4E). This indicated that WASH was an essential actin regulator of endocytosis.

Next, we investigated whether WASH was activated by ubiquitylation of residue K220, as previously described for its endosomal function (Hao et al., 2013). However, a WASH K220R mutant incapable of ubiquitylation rescued loss of WASH to the same extent than wildtype WASH (Figure 4F) indicating that ubiquitylation of WASH was dispensable for WASH activation during endocytosis. In contrast, loss of function mutation of a phosphorylation site at Y141 previously implicated in WASH activity in natural killer (NK) cells (Huang et al., 2016) failed to rescue loss of WASH, whereas the gain of function mutation Y141E rescued similar to wildtype WASH (Figure 4G). Again, this data distinguished WASH regulation during endocytosis from its typical function on endosomes.

### WASH acted during late stages of endocytosis

We then addressed the role of WASH during HPV16 endocytosis more specifically. Unaffected HPV16 binding to WASH KO cells suggested that recycling and binding receptor presentation were in principle intact (Figures S4D and S4E). Moreover, plasma membrane presentation of the internalization receptor candidates CD151 and EGFR were largely unaffected (Figure S4C), so we investigated how loss of WASH may interfere with endocytosis.

Analysis of virus-containing pits in ultra-thin section transmission EM revealed that endocytic pits were morphologically unaltered in WASH KO compared to wild type cells, i.e., they were fully formed and partially constricted at the neck (Figure 5D). Thus, early stages of endocytic vesicle formation such as induction and membrane invagination were independent of WASH. Strikingly, the average number of virus-containing plasma membrane invaginations more than doubled in the absence of WASH (Figures 5D). Hence, endocytosis was stalled at a late stage such as scission in WASH KO cells. Why, however, did loss of WASH fail to replicate the phenotype of globally perturbed actin polymerization, where virus-filled tubules formed (compare Figure 1)? We reasoned that since cytochalasin D interfered not only with WASH-mediated but also with cortical actin polymerization cortical actin may restrict membrane feeding into endocytic pits. In this case, cytochalasin D treatment in WASH knockout cells should replicate the earlier result. And in fact, virus-filled tubules arose also in the case of WASH knockout cells when treated with cytochalasin D (Figure 5E). Taken together, the roles of actin polymerization in vesicle scission, of WASH as the single NPF to promote actin polymerization, and the loss of WASH stalling endocytosis at a late stage of vesicle formation strongly indicated that WASH regulated endocytic vesicle scission but not endocytic pit formation.

**Figure 5.**
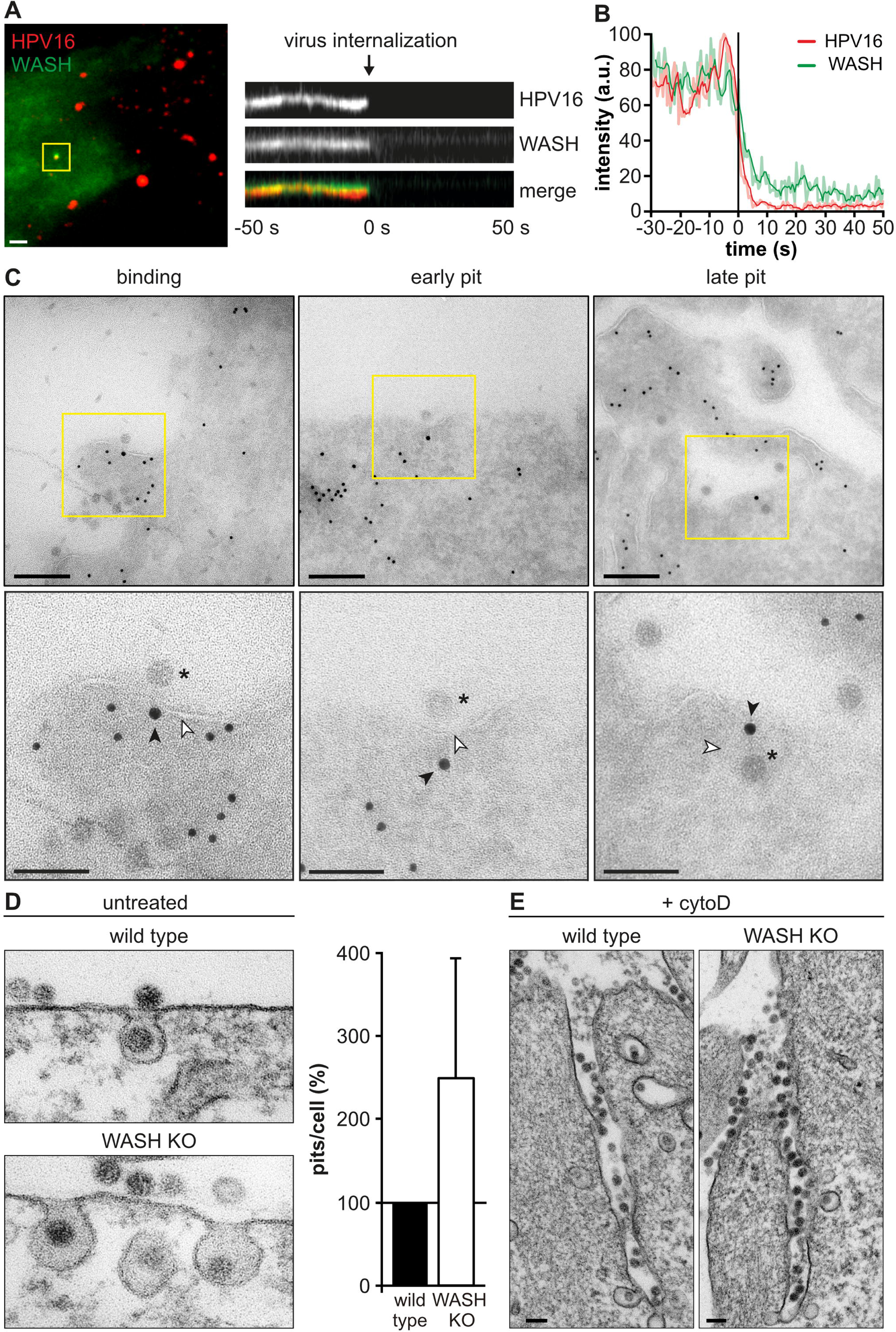
WASH associates early, but acts late in endocytosis. (A) HeLa ATCC cells transfected with EGFP-WASH were infected with HPV16-AF647. Cells were imaged by live cell TIRF-M at 1 h p.i.. Movies were acquired with 0.5 Hz frame rate for 5 min. HPV16 entry events were identified manually after background subtraction and filtering. Shown is a kymograph of the virus entry event highlighted by the yellow box, and the corresponding EGFP-WASH signal. Scale bar is 2 µm. (B) Intensity profiles of HPV16 and WASH (light red/green) as well as moving averages (intense red/green) of the virus entry event shown in (A) depicted relative to the half time of virus loss (t = 0). (C) HeLa ATCC cells were transfected with EGFP-WASH and infected with HPV16. At 6 h p.i., cells were subjected to immunogold labeling of GFP (WASH, 15 nm gold) and actin (10 nm gold) on ultra-thin cryosections analyzed by transmission EM. Asterisks indicate HPV16 particles, black and white arrowheads indicate WASH staining and the membrane, respectively. Scale bars are 200 nm and 100 nm for overviews and insets, respectively. (D) NIH-3T3 wild type and WASH KO cells infected with HPV16-GFP were subjected to ultra-thin section transmission EM at 6 h p.i.. The number of virus-filled plasma membrane invaginations was determined for 43 cells per cell line in two independent experiments. Total pit numbers were normalized to wild type cells and are depicted as mean ± SD. Scale bars are 100 nm. (E) Infection of NIH-3T3 wild type and WASH KO cells was carried out in presence of 5 µg/ml cytoD. At 6 h p.i., cells were processed for ultra-thin section transmission EM. Scale bars are 100 nm.

To directly stimulate vesicle scission, WASH would have to be present at endocytic sites. Recruitment of WASH to specialized TEMs marked by CD151 (Scheffer *et al*., 2013) was probed by a proximity ligation assay (PLA) (Söderberg et al., 2006). A small fraction of virus particles co-localized with the PLA signal of WASH and CD151 (Figure S6A). Hence, WASH may indeed act at sites of cargo uptake. Only a limited association of HPV16 with WASH/CD151 structures occurred within coincidence detection limits in fixed samples. This was expected, since virus uptake occurs asynchronously over many hours, so that only few viruses interact with specialized TEMs at any given time (Schelhaas *et al*., 2012; Becker *et al*., 2018). However, to more conclusively demonstrate recruitment of WASH to endocytic sites and to gain information on recruitment dynamics, WASH association in relation to cargo uptake was analyzed by live cell TIRF-M. WASH was detected together with HPV16 for more than 50 seconds prior to uptake and co-internalized with the cargo (Figures 5A and 5B, Suppl. Movie 3). To pinpoint the stage during which WASH is recruited to endocytic sites, immunogold labelling and EM was performed. Indicative of specific labeling, WASH was found on endosomes (Figure S6B). Moreover, it was observed close to virus particles associated with flat plasma membrane regions (binding), slightly curved membranes (early pit), and with fully formed endocytic pits (late pit) (Figure 5C). In fact, quantification of the immunogold EM revealed that WASH signals at the plasma membrane were similarly specific as WASH labeling on endosomes Figure S6C). In conclusion, WASH was recruited to the plasma membrane already at a very initial stage of pit formation, and remained during all stages of pit maturation (Figure 5C). Thus, WASH was recruited already early during vesicle formation, although it likely exerted its NPF function only during scission similar to its function on endosomes (Derivery *et al*., 2009; Gomez and Billadeau, 2009).

## Discussion

Here, we elucidate mechanistic details of the endocytic process by which HPV16 is internalized into cells. Originally distinguished from other modes of endocytosis predominantly in negative terms, e.g., clathrin-, caveolin-, dynamin-, cholesterol-independent and morphologically distinct from macropinocytosis, this work now provides evidence for the existence of a WASH-mediated scission mechanism during clathrin-independent endocytosis. Cargo is internalized by inward budding of the plasma membrane in distinguishable steps. Thus, vesicles are most likely formed by *de novo* assembly of endocytic machinery in a modular manner, a mode comparable to CME and distinct from macropinocytosis (Figure 6). Our results implicated WASH in the stimulation of branched actin polymerization most likely for scission. The role of WASH in virus uptake was clearly distinguishable from its role in retrograde trafficking due to its independence of retromer-associated sorting complexes, ubiquitylation at K220, and members of the WASH complex (SHRC). Hence, our work describes an unexpected direct function of WASH in endocytosis. Since WASH has not been attributed to any other endocytic pathway, it defines the molecular identity of this endocytic process.

**Figure 6.**
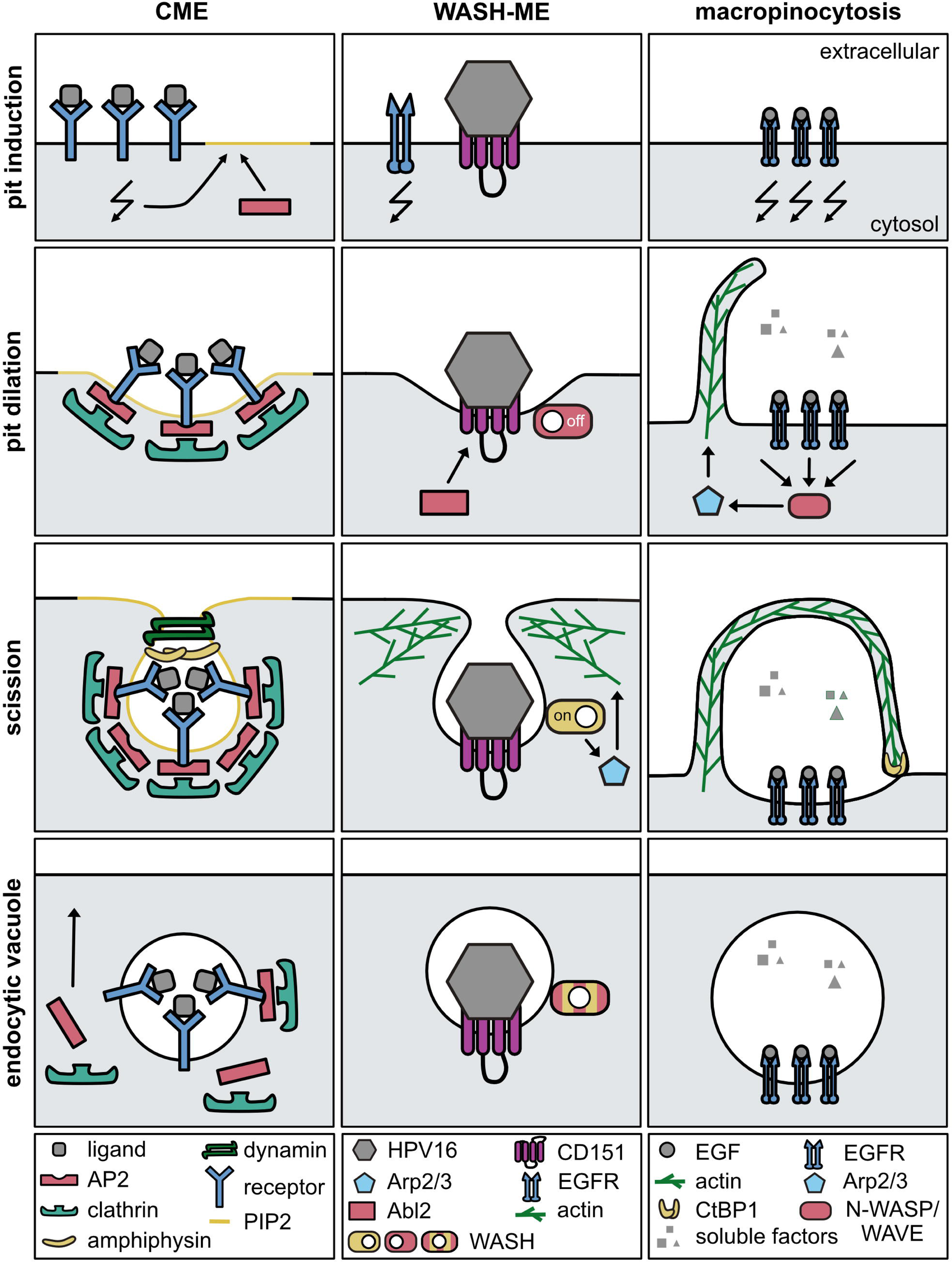
Model. Schematic model of the mechanistic regulation of endocytic vesicle formation during WASH-ME in comparison to CME and macropinocytosis. Additional regulators involved in the latter mechanisms were omitted for clarity.

The endocytic landscape includes a variety of processes, where endocytic vesicle formation is achieved by different machineries for different purposes. These typically diverge in morphology, key cargos, and molecular features (Doherty and McMahon, 2009). Often, the pinocytic mechanisms are subdivided into CME and clathrin-independent endocytosis, the latter of which applies to HPV uptake. Clathrin-independent endocytosis includes mostly mechanisms requiring cholesterol and glycosphingolipids, and the completion of vesicle formation by a scission event often depends on dynamin. Since HPV endocytosis is independent of both cholesterol and dynamin (Schelhaas et al., 2008, Schelhaas et al., 2012, Spoden et al., 2008, Spoden et al., 2013), it likely uses separate machineries.

Generally, scission may occur by one of several hypothesized mechanisms involving e.g. mechanoenzymatic scission by dynamin, membrane-lipid-reorganizing proteins or complexes (actin cytoskeleton and CtBP/BARS), or the appearance of a lipid diffusion barrier combined with a pulling force induced by molecular motors (friction-driven fission) (summarized in Renard et al., 2018). As mentioned, scission is mediated by the large GTPase dynamin for many endocytic mechanisms. In mechanoenzymatic scission, e.g. during CME, dynamin polymerizes to form a collar that compresses the vesicle neck by an extensively discussed mechanism leading to membrane fission (Hinshaw and Schmid, 1995; Takei et al., 1995; Sweitzer and Hinshaw, 1998; Morlot and Roux, 2013). In contrast, HPV uptake occurs independently of dynamin (Spoden *et al*., 2008; Schelhaas *et al*., 2012). HPV uptake involves actin polymerization, which coincided with vesicle scission in a timely fashion that resembled dynamin recruitment during CME (Merrifield *et al*., 2002; Merrifield, 2004). Hence, actin polymerization likely facilitates vesicle scission by a force-driven mechanism rather than as anchoring structure for other scission factors. Alternative roles for actin polymerization in scission appear less likely including membrane-lipid-reorganization followed by line tension-dependent scission as described for Shiga toxin (Römer et al. 2010) or as lipid diffusion barrier (friction-driven fission) as proposed by Simunovic and colleagues (2017): The first typically involves cholesterol-rich membranes and the second molecular motors such dynein or myosin II, whereas HPV uptake requires neither (Schelhaas et al., 2008, Schelhaas et al., 2012, Spoden et al., 2008, Spoden et al., 2013). Moreover, a hallmark of line tension dependent scission and friction-driven fission is formation of tubular invaginations, which were absent upon WASH knockout. While this does not exclude that such mechanisms contribute to the fidelity of scission particularly in light of cortical actin polymerization, it renders a force-driven scission mechanism through WASH-promoted actin polymerization more likely. How actin polymerization creates the force for vesicle scission remains elusive. However, in analogy to observations from CME or endocytosis in *Xenopus* oocytes, actin polymerization towards the vesicle neck may serve to constrict and propel the vesicle away from the plasma membrane (Bement et al., 2003; Sokac et al., 2003; Collins et al., 2011).

How WASH would be recruited to the plasma membrane, remains elusive. WASH recruitment to endosomes is regulated by incorporation into the WASH regulatory complex, similar to SNX2 recruitment as a heterodimeric complex with SNX5 or 6 (Wassmer et al., 2007; Derivery *et al*., 2009; Wassmer et al., 2009; Jia et al., 2010). However, since WASH can act independently of the SHRC for HPV endocytosis, and since SNX-BARs are dispensable in this context, WASH recruitment is likely to be different. For WASH, recruitment may be facilitated by FAM21 through direct phospholipid binding, as *in vitro* studies revealed FAM21 binding to a variety of phospholipids (Derivery, Sousa, *et al*., 2009; Jia *et al*., 2010).

Dissimilar to WASH activation at endosomes (Hao *et al*., 2013), ubiquitination of WASH was not essential for HPV endocytosis, whereas an ill-defined gain of function of WASH mediated by phosphorylation of Y141 that has been previously described in NK cells (Huang *et al*., 2016) may be important in this context. In NK cells, phosphorylation of WASH is mediated by lymphocyte-specific tyrosine kinase (Lck), which is not expressed in fibroblasts or epithelial cells. Instead, perhaps the related Abl2 kinase, previously implicated in HPV endocytosis may function instead to activate WASH by phosphorylation of Y141 (Bannach et al., 2020).

How HPV uptake is induced by ligand-receptor interactions is an important question. Likely, this occurs only during specific cell states, as WASH is infrequently observed at the plasma membrane. Evolutionary, it is most probable that HPV16 exploits a cellular process that is easily inducible or already active upon infection of target cells. This situation exists upon epidermal wounding, where basal keratinocytes of the mucosal epidermis as primary targets become accessible to the virus (Doorbar, 2005; Roberts et al., 2007; Aksoy et al., 2017). Wounding and other cellular responses induce epithelial to mesenchymal transition and cell migration, which are accompanied by the remodeling of cell-matrix adhesion complexes, such as focal adhesions and hemidesmosomes (HDs) (Jones et al., 1998; Borradori and Sonnenberg, 1999; Webb et al., 2002; Ezratty et al., 2005; Walko et al., 2015). To date, relatively little is known about the dynamics of HD remodeling during cell migration, but HD containing plasma membrane domains are rapidly endocytosed upon detachment from the underlying extracellular matrix (ECM) (Owaribe et al., 1990). Since HDs contain integrin α6 and CD151, which are part of the specialized TEMs that mediate HPV16 endocytosis (Scheffer *et al*., 2013; Walko *et al*., 2015; Mikuličić et al., 2019), The endocytic machinery facilitating HPV uptake may also promote HD uptake during wound healing to aid cell migration and thereby wound closure. In line with this notion, CD151 TEMs co-internalize with the virus and are remodeled upon growth factor signaling (Scheffer *et al*., 2013; Mikuličić *et al*., 2019). Interestingly, a CME sorting motif in the cytosolic domain of CD151 is not required for its function during HPV endocytosis supporting its specialized role for retrieval of functionalized TEMs (Liu et al., 2007; Scheffer *et al*., 2013). Another essential process during wound healing is matrix remodeling. Thus, it is noteworthy that WASH was previously implicated in apical endocytosis of extracellular material in the *Drosophila* airway epithelium during embryogenesis (Tsarouhas et al., 2019). Hence, HPV endocytosis may also contribute to matrix remodeling as a specialized mechanism for internalization of extracellular material.

While future research will have to address the cellular role of HPV endocytosis in more detail, its role in pathogen invasion may be of considerable interest as well. For instance, WASH is recruited to the plasma membrane during *Salmonella* infection presumably serving as one of several entry pathways (Hänisch et al., 2010). Moreover, WASH regulatory complex protein FAM21 co-localizes with a subset of VV particles in plasma membrane lipid rafts (Huang et al., 2008). Thus, HPV endocytosis may constitute an important entry route for pathogens. Accordingly, viruses such as IAV and LCMV exploit pathways with endocytic vacuole morphology and mechanistic requirements similar yet not identical to HPV16 endocytosis (Sieczkarski and Whittaker, 2005; Quirin *et al*., 2008; Rojek et al., 2008; de Vries et al., 2011).

## Acknowledgements

We thank I. Fels and D. Kaiser (Institute of Cellular Virology, Münster, Germany) for technical support and U. Westerkamp and M. Dominguez (Institute of Cellular Virology, Münster, Germany) for assistance with experiments. We are indebted to numerous individuals for sharing valuable reagents as indicated in the STAR methods. We acknowledge the Infectious Diseases Imaging Platform at the University Hospital Heidelberg, K. Richter from the Core Facility Unit Electron Microscopy at the DKFZ Heidelberg and S. Hillmer from the Electron Microscopy Core Facility of the University Heidelberg for their technical support. This work was supported by funding of the German Research Foundation (DFG) to MS (grants SCHE 1552/6-2 and INST211/817A09), and to SB (grant BO 4340/2-1). T.E.B.S. acknowledges support by the Helmholtz Society through HGF impulse fund W2/W3-066.

## Author contributions

Conceived and designed experiments: PB, LK, PSV, CB, TS, MS; Performed experiments: PB, LK, PSV, LG, CB, DB, JK; Analyzed data: PB, LK, PSV, LG, CB, DB, JK, SB, TS, MS; Resources: PD, SB, TS, MS; Writing: PB, LK, MS with input from the other authors.

## STAR methods

### KEY RESOURCES TABLE

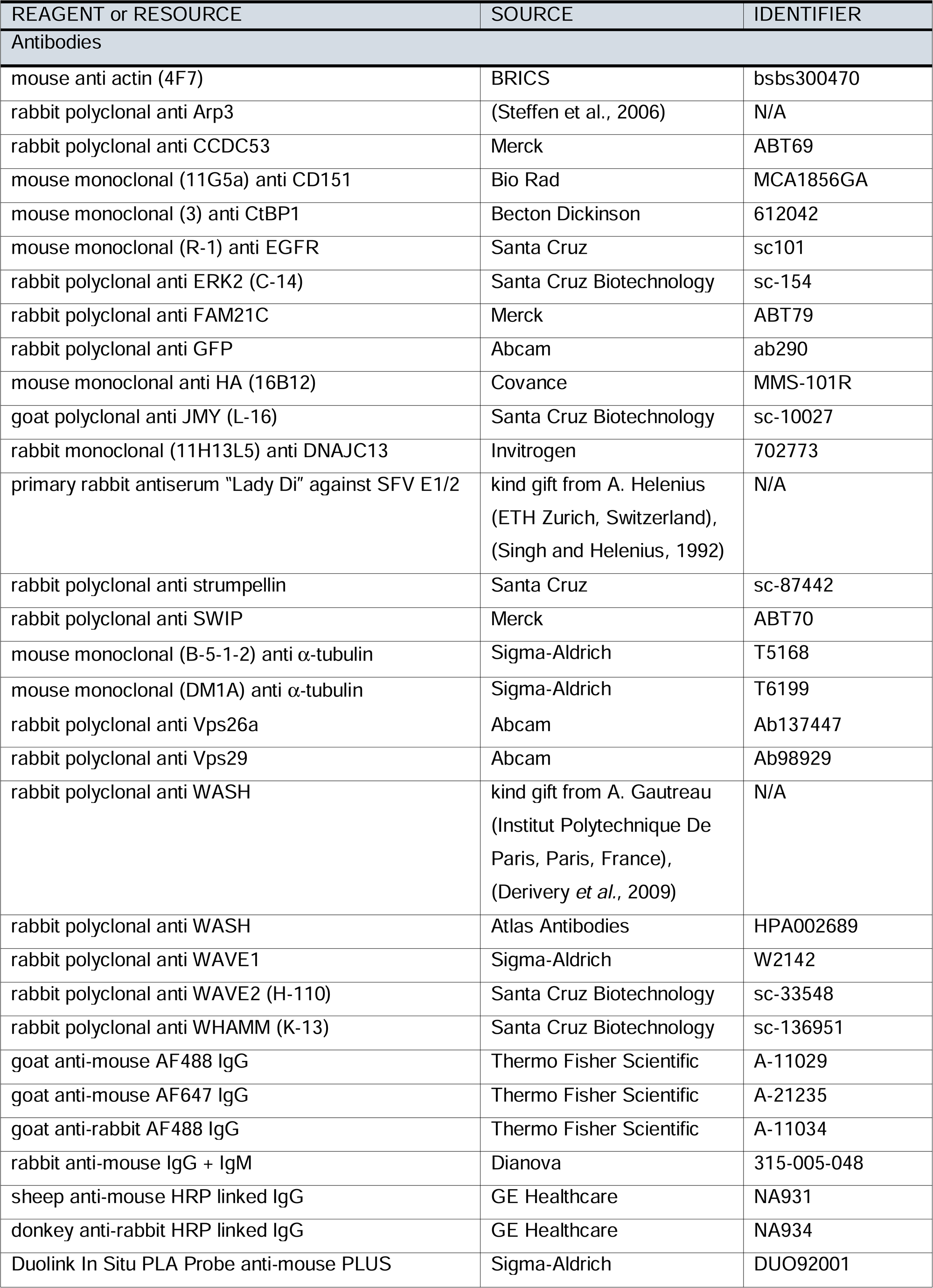
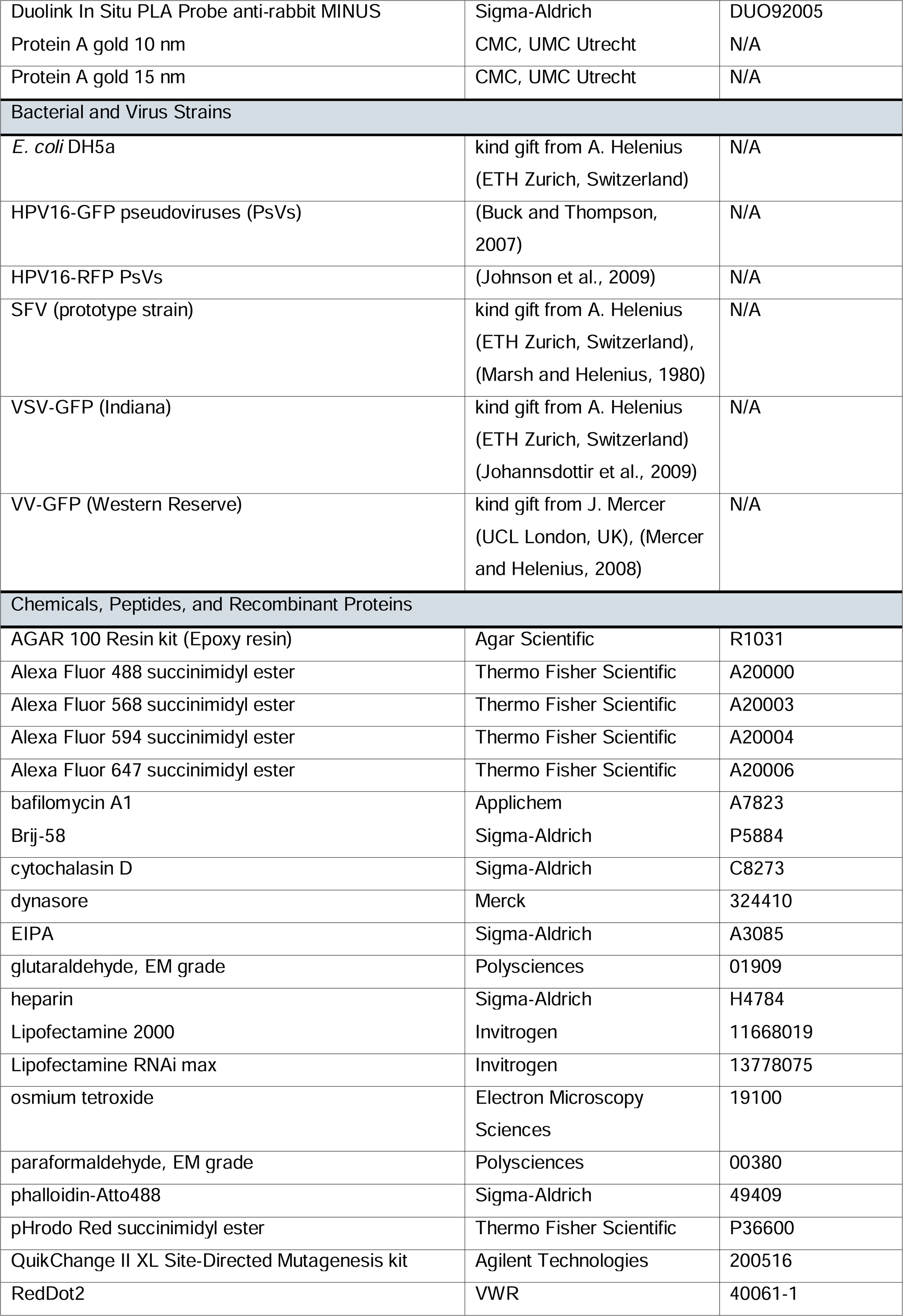
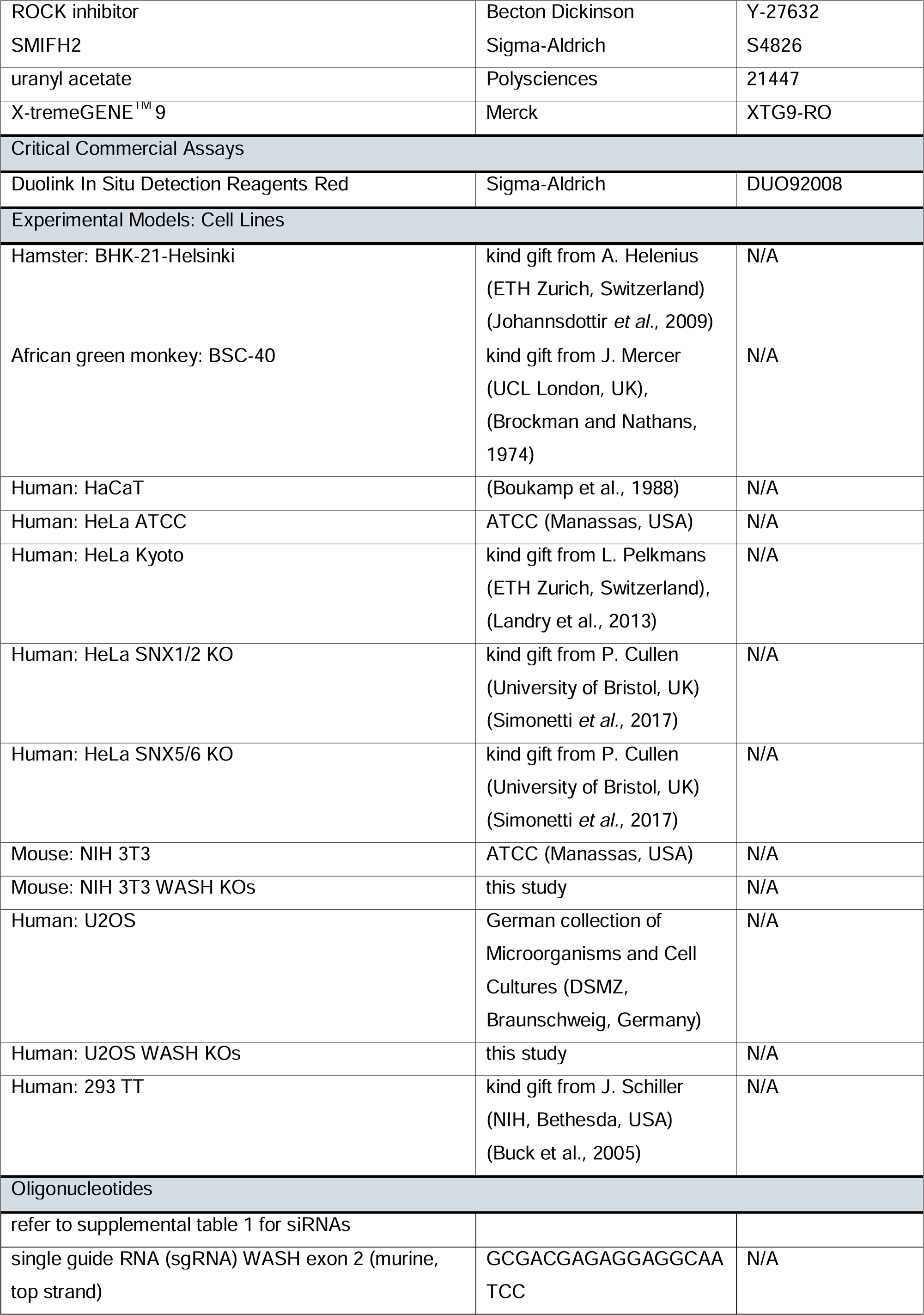
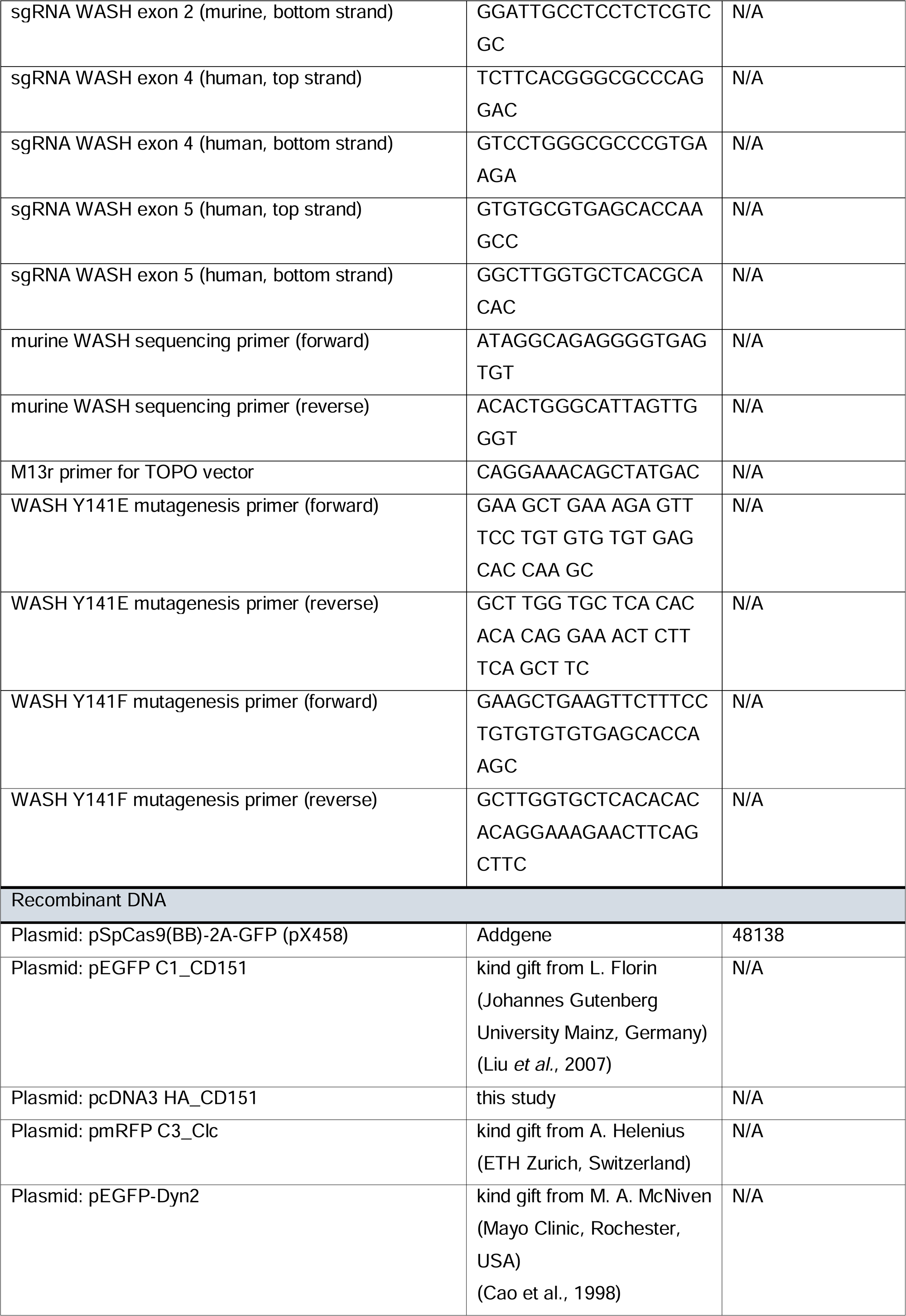
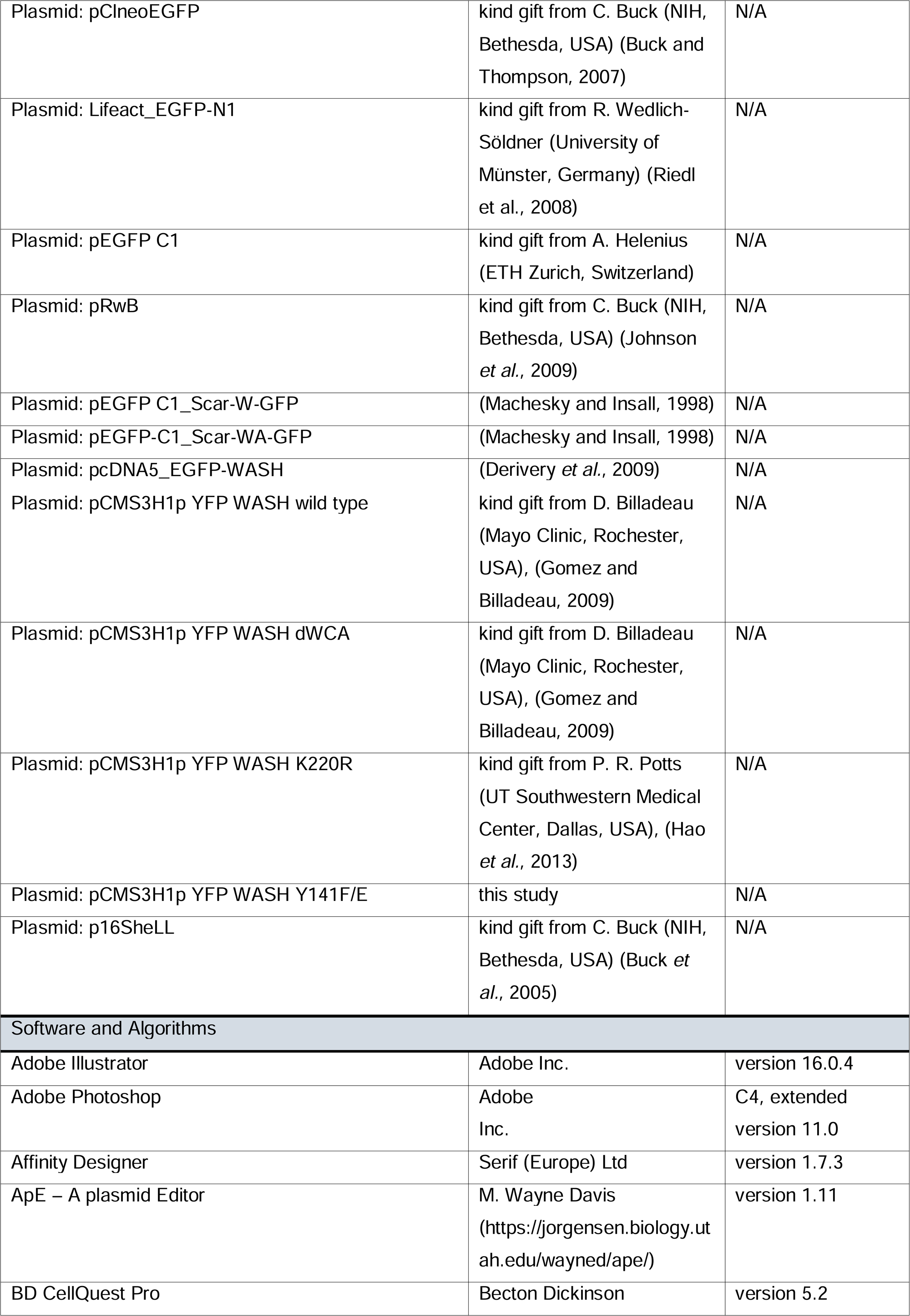
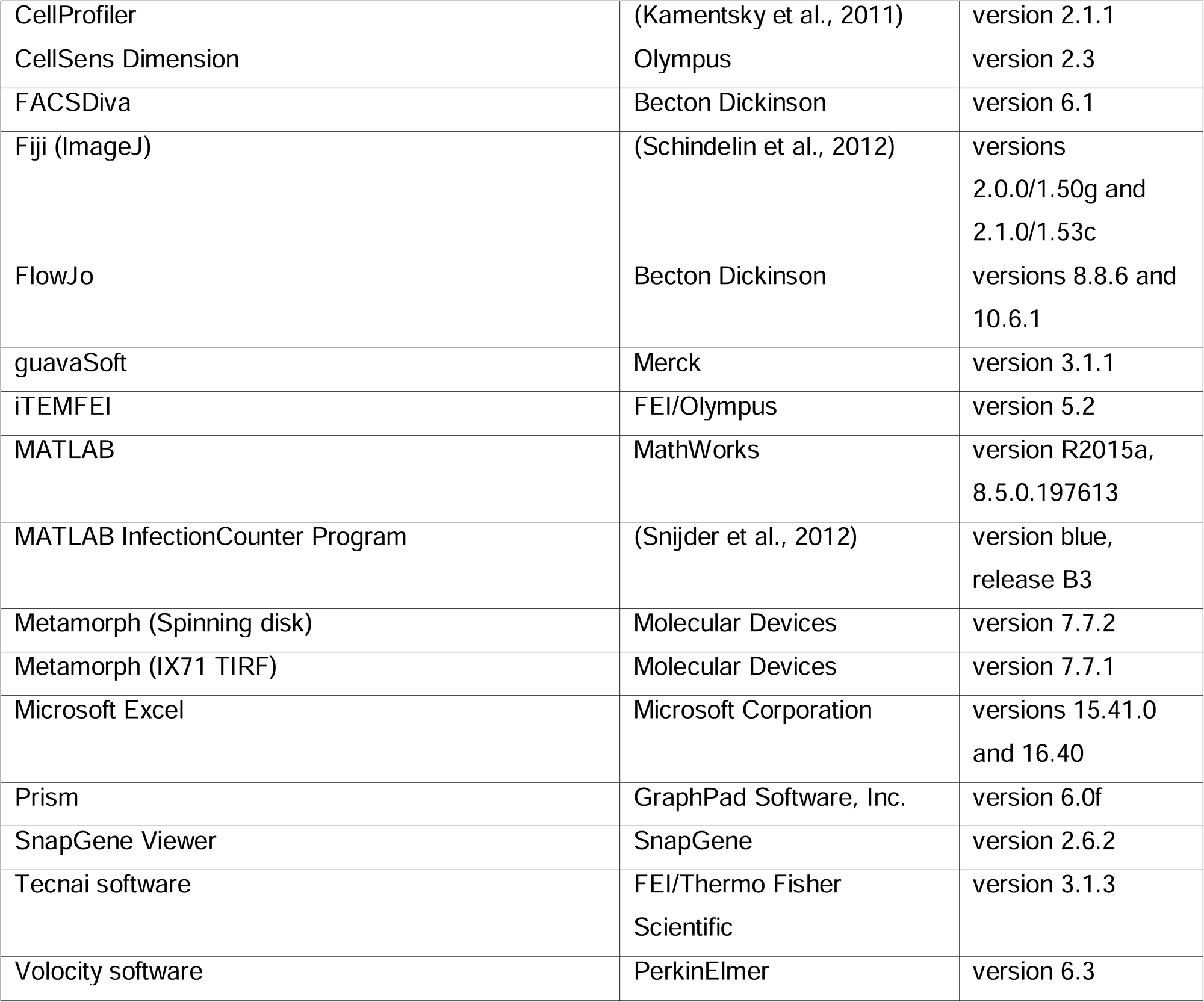

### RESOURCE AVAILABILITY

#### Lead Contact

Further information and requests for resources should be directed and will be fulfilled by the lead contact, Mario Schelhaas.

#### Materials Availability

Plasmids generated in this study are available from the lead contact. WASH knock out cell lines described in this study are available from Theresia Stradal with a completed Material Transfer Agreement.

#### Data and Code Availability

All data reported in this paper will be shared by the lead contact upon request. The paper does not report original code. Any additional information required to reanalyze the data reported in this paper is available from the lead contact upon request.

### EXPERIMENTAL MODEL AND SUBJECT DETAILS

#### Cell lines

HeLa ATCC, HeLa Kyoto, HeLa SNX1/2 KO, HeLa SNX5/6 KO (all female origin) and HaCaT cells (male origin) were cultured in high glucose Dulbecco’s modified eagle medium (DMEM, D5796 Sigma-Aldrich) supplemented with 10% fetal bovine serum (FBS). 293TT cells (female origin) were grown in DMEM with 10% FBS and 400 µg/µl hygromycin B. U2OS wild type and WASH KO cells (female origin) as well as murine NIH 3T3 wild type and WASH KO cells (male origin) were cultured in DMEM supplemented with 10% FBS and 1% non-essential amino acids (NEAA). Primate, non-human BSC-40 cells (sex unspecified) were grown in DMEM supplemented with 10% FBS, 5% NEAA and 5% sodium pyruvate. Hamster BHK-21-Helsinki cells (male origin) were cultured in Glasgow’s minimum essential medium (GMEM) supplemented with 10% FBS. All cells were cultivated in a humidified atmosphere at 37°C and 5% CO_2_ and routinely tested for mycoplasma contamination.

#### Bacteria strains

Chemocompetent *E. coli* DH5a used for plasmid preparation were grown in lysogeny broth (LB) medium supplemented with antibiotics at 37°C and 210 rpm.

#### Virus strains

VV-GFP (Western Reserve) containing a fluorescent version of the core protein A5 was propagated and titrated on BSC-40 cells in standard growth medium and purified as described previously (Mercer and Helenius, 2008). SFV (prototype strain) was propagated and titrated on BHK-21-Helsinki cells as previously described (Marsh *et al*., 1984). Infection was carried out in GMEM supplemented with 10% FBS and 10% Tryptose Broth. VSV-GFP (Indiana) expressing an additional transgene encoding GFP was propagated on BHK-21-Helsinki cells grown in GMEM supplemented with 10 mM HEPES (pH 6.5). At 1 h p.i., GMEM supplemented with 30 mM HEPES (pH 7.3) and 10% Tryptose Phosphate Broth and 1% FBS was added, as previously described (Johannsdottir *et al*., 2009). The virus was titrated on BHK-21-Helsinki cells grown in RPMI supplemented with 30 mM HEPES (pH 6.5). HPV16 PsVs were produced as described in the method details.

### METHOD DETAILS

#### Cloning and plasmid purification

##### Cloning of HA-CD151 construct

Lentiviral expression constructs of HA-tagged CD151 were a kind gift from M. Hemler (Dana Farber Cancer Institute and Harvard Medical School, Boston, USA) (Hwang et al., 2019). For subcloning, HA-CD151 was cut from the lentiviral vector using *EcoR*I and *Xba*I for 2 h at 37°C. A pcDNA3 expression vector was cut using the same enzymes and conditions. For purification, both samples were run on a 2% acrylamide gel. The DNA was visualized with ethidium bromide and bands representing the cut pcDNA3 backbone as well as HA-CD151 (insert) were isolated from the gel. Gel extraction was performed with a NucleoSpin Gel and PCR Clean-up kit (Macherey-Nagel). DNA concentrations were determined with help of a 1.5% agarose gel by comparison to the marker bands (Gene Ruler 1 kb DNA ladder, Thermo Scientific). Insert and backbone were ligated by incubation with the T4 ligase overnight at 16°C.

##### Plasmid purification

Chemocompetent *E. coli* DH5a were incubated with 5 µl ligation product for 10 min on ice. Heat shock was performed by 60-90 sec incubation at 42°C. Immediately afterwards, bacteria were incubated on ice for 5 min before LB medium was added. Bacteria were grown at 37°C and 350 rpm for 30-60 min and plated on LB agar plates with antibiotics using inoculation loops. Plates were incubated overnight at 37°C and inspected for colony growth the next day. For HA-CD151 cloning, several overnight cultures with LB supplemented with antibiotics were inoculated with one colony each. Plasmids were purified using the NucleoSpin Plasmid kit (Macherey-Nagel) and sent for sequencing by Eurofins Genomics (Luxembourg). Sequence analysis was performed with ApE. For use in experiments, plasmids were purified using the NucleoBond Xtra Maxi kit (Macherey-Nagel).

##### Site Directed Mutagenesis

WASH Y141 mutants were generated following the protocol of the QuikChange II XL Site-Directed Mutagenesis kit (Agilent Technologies). Primers were designed based on the manufacturers guidelines and are listed in the key resources table. Following the PCR and DpnI digest, chemocompetent *E. coli* DH5a were transformed and plasmids were purified as described above. Successful mutagenesis was controlled by PCR.

##### sgRNA design and cloning

sgRNAs were provided by a CRISPR design tool (CRIPR.mit.edu or CCTop). Specifically, the murine gene was disrupted by sgRNAs targeting exon 2 and the human gene by simultaneously targeting exons 4 and 5 (key resources table). Top and bottom strands of sgRNAs were annealed using T4 ligase for 30 min at 37°C and cloned into pSpCas9(BB)-2A-GFP (pX458) by digestion with *Bbs*I and ligation with T4 ligase (6 cycles: 37°C for 5 min, 21°C for 5 min). Residual linearized DNA was removed by treatment with PlasmidSafe exonuclease at 37°C for 30 min. Chemically competent *E.coli* DH5a were transformed with the ligation product as described above.

#### Generation and characterization of WASH KO cell lines with CRISPR/Cas9

NIH-3T3 and U2OS cells were plated in 6-well plates and maintained in DMEM (4.5 g/L glucose, Invitrogen, Germany) supplemented with 10% (v/v) FBS (Sigma, Germany), 1 mM sodium pyruvate, 1x non-essential amino acids and 2 mM L-glutamine at 37°C in a humidified 7.5% CO2-atmosphere overnight. Cells were genome edited using the CRISPR/Cas9 technology (Ran et al., 2013) to generate WASH KO cell lines. Selected sgRNAs (key resources table) were cloned into pX458 allowing simultaneous expression of sgRNA, Cas9 and selection via EGFP expression as described above. The resulting constructs were transfected into NIH-3T3 or U2OS cells, respectively. Plasmids (1 µg) and X-tremeGene^TM^ 9 (3 µl) were diluted in 100 µl optiMEM, incubated for 30 min at room temperature and added to cells for 16-24 h. Transfection efficiency was monitored using an EVOS® FL Cell Imaging System (Thermo Fisher, Germany). Cells were grown to confluence and subsequently single, GFP-positive cells were sorted into 96-well plates by flow cytometry using a FACSAria II instrument (BD Biosciences) and FACSDiva software. Sorted cells were maintained in growth medium supplemented with penicillin (50 Units/ml)/streptomycin (50 mg/ml) (Thermo Fisher Scientific) and containing 30% conditioned medium and 10 µM ROCK inhibitor (BD Biosciences). After approximately 10 days, clones were picked from single wells and expanded. Derived cell clones were screened for the absence of WASH expression by Western blotting and NIH-3T3 clones lacking detectable amounts of WASH were subjected to genomic sequencing as described (Kage et al., 2017). Cells from confluent 6 cm dishes were trypsinized, pelleted and lysed by overnight incubation in lysis buffer (100 mM Tris pH 8.5, 5 mM EDTA, 0.2% SDS, 200 mM NaCl, 20 mg/ml proteinase K) at 55°C. Nucleic acid extraction was performed by ethanol precipitation. Addition of 700 µl 100% ice cold ethanol was followed by centrifugation at 16,000 x g at 4°C for 30 min. The pellet was washed with 400 µl 70% ice-cold ethanol and samples were dried at 45°C for 20 min. DNA was dissolved in 100 µl deionized water at 4°C overnight and served as template in PCR using GoTaq G2 flexi DNA polymerase. The selected primer pair (key resources table) revealed a PCR product of 330 base pairs. PCR products were examined on 2% agarose gels and appropriate samples purified with a NucleoSpin Gel and PCR clean-up kit (Macherey-Nagel). DNA fragments were cloned into a zero blunt TOPO vector (Zero Blunt TOPO Cloning Kit for Sequencing, Invitrogen) for 5 min at room temperature. After transformation as above, single bacterial colonies were inoculated overnight and plasmid DNA purified using NucleoSpin Plasmid kit (Macherey-Nagel). Sequencing of isolated plasmid DNA was carried out by MWG-Biotech using a standard M13r sequencing primer (key resources table). Clones were examined for frameshift mutations and mono- or biallelic deletions/insertions using SnapGene Viewer software. Mutations or deletions generating stop codons shortly downstream of the target site were defined as ‘null’ alleles. Cell populations exclusively harboring such alleles out of more than 50 sequencing reactions were selected for further analyses. For WASH targeted clones of murine NIH-3T3 cells, three clones were identified that did not display a wild type allele in more than ten sequencing reactions. All clones showed similar effects on HPV16 infection, thus only one clone is shown in this study. For WASH targeted clones of human U2OS cells no clear sequencing result was obtained probably due to pseudogenes being targeted by sequencing primers.

#### Western blotting

For analysis of protein amounts after siRNA treatment or CRISPR/Cas9 mediated KO, lysates were prepared in 2x sample buffer (4% SDS, 20% glycerol, 0.01% bromophenol blue, 100 mM Tris HCl (pH 6.8), 200 mM DTT). Samples were denatured for 5 min at 95°C and loaded on polyacrylamide gels. For stacking gels, 5% polyacrylamide was used and separating cells were 6 or 8% for JMY and WHAMM, respectively. All other proteins were separated on 10% gels. Electrophoresis was performed in Laemmli running buffer (0.1% SDS, 25 mM Tris, 192 mM glycine). Proteins were transferred from gels to nitrocellulose membranes for 50 min at 400 mA in pre-cooled transfer buffer (192 mM glycine, 25 mM Tris, 10% methanol). After transfer, membranes were blocked in 5% milk powder (MP) in Tris-buffered saline (TBS) supplemented with 0.2% Tween 20 (TBS-TMP) or in 0.2-3% bovine serum albumin (BSA) for at least 30 min. Primary antibodies were diluted in TBS-TMP or BSA and membranes were incubated for 2 h at room temperature or overnight at 4°C. Three washes with TBS supplemented with Tween 20 (TBS-T) were followed by incubation with anti-mouse or -ra bbit secondary antibodies conjugated to HRP diluted in TBS-TMP or BSA. Membranes were washed twice with TBS-T and once with TBS before the signal from HRP-conjugated antibodies was revealed using enhanced chemiluminescence (ECL) or ECL prime and photographic films or a chemiluminescence imager (ChemoStar Touch, Intas).

#### Virus production

##### Production of HPV16 PsVs

PsV production was performed as previously described (Buck *et al*., 2005). A total of 1.8 x 10^7^ 293TT cells were seeded in 145 mm cell culture dishes. The next day, cells were co-transfected with p16sheLL and the reporter plasmid pClneoEGFP (GFP) or pRwB (RFP). Both plasmids (30 µg each) as well as Lipofectamine 2000 (132.5 µl) were diluted in optiMEM and incubated for 5 min at room temperature. The DNA dilution was added to the Lipofectamine 2000 dilution and samples were incubated for 20 min at room temperature before the transfection mix was added to fresh growth medium in the dishes. At 48 h post transfection, cells were harvested and pelleted. For cell lysis and virus maturation, the pellet was incubated with 0.35% Brij 58, 0.2% Plasmid Safe DNase and 0.2% benzonase for 24 h at 37°C on an overhead rotator. PsVs were purified using a linear 25%-39% OptiPrep density gradient. A PsV fraction at around 30% OptiPrep was collected with a needle and analyzed for virus content and purity by Coomassie staining of SDS-PAGE gels.

##### Labeling of HPV16 PsVs

HPV16 PsVs were incubated with Alexa Fluor 488, 568, 594 or 647 succinimidyl ester in virion buffer (635 mM NaCl, 0.9 mM CaCl_2_, 0.5 mM MgCl_2_, 2.1 mM KCl in PBS, pH 7.6) using a 1:8 molar ratio of L1 to the dye for 1 h on an overhead rotator (Schelhaas *et al*., 2008; Samperio Ventayol and Schelhaas, 2015). Free dye was removed by ultracentrifugation using a 15-25-39% OptiPrep step gradient. The labeled virus between the 25% and 39% OptiPrep fraction was collected with a needle. The PsV concentration was determined by SDS-PAGE and subsequent Coomassie staining. The labeled virus was characterized by binding to glass coverslips and HeLa ATCC cells (Samperio Ventayol and Schelhaas, 2015). Labeling of PsVs with pHrodo was achieved following the same protocol. Virus characterization was performed in citric acid buffer (pH 4.4) (Samperio Ventayol and Schelhaas, 2015).

#### Infection experiments

##### Infection of KO cells

NIH-3T3 (4 x 10^5^ cells/well) and U2OS (both 5 x 10^4^ cells/well) wild type and WASH KO cells were seeded in 12-well plates. The next day, the growth medium was replaced with 300 µl fresh growth medium and HPV16-GFP was added to result in 20% infection in wild type cells. The virus was bound on a shaker at 37°C. At 2 h p.i., the inoculum was replaced by fresh growth medium and infection was continued. At 48 h p.i., cells were trypsinized and fixed in 4% PFA for 15 min at room temperature. Cells were resuspended in FACS buffer (250 mM EDTA, 2% FBS, 0.02% NaN_3_ in PBS) and analyzed for infection (percentage of GFP positive cells) by flow cytometry (FACSCalibur, Becton Dickinson). Gating of infected cells was done with help of uninfected controls. The percentage of infected cells was normalized to the respective wild type cells using Microsoft Excel.

##### Inhibitor studies

HeLa ATCC were seeded in 12-well plates (5 x 10^4^ cells/well) about 16 h prior to experimentation. For HPV16 infection experiments, small compound inhibitors and solvent controls were diluted in growth medium, while they were diluted in infection medium (RPMI supplemented with 30 mM HEPES, pH 6.5) for VSV infection. Cells were pre-treated with inhibitors or solvent controls for 30 min at indicated concentrations and infected with HPV16-GFP as described above. The inoculum was replaced at 2 h p.i. and infection was continued in presence of the inhibitor. At 12 h p.i., inhibitors were exchanged for 10 mM NH_4_Cl/10 mM HEPES in growth medium to reduce cytotoxicity (Schelhaas *et al*., 2012). Cells were fixed and processed for flow cytometry analysis as described above. For infection with VSV-GFP, the virus was added to the infection medium +/-inhibitor to result in 20% infection in solvent treated controls. VSV-GFP was bound for 2 h on a shaker at 37°C until the inoculum was replaced with 1 ml growth medium. Cells were trypsinized and fixed at 6 h p.i. as described for HPV16.

##### Infection studies after siRNA-mediated depletion

For siRNA-mediated knockdown, 2 x 10^3^ or 2 x 10^4^ HeLa Kyoto cells were reverse transfected in 96-well optical bottom plates or 12-well plates, respectively. Transfection was performed using 0.2 µl (96-well) or 0.5 µl (12-well) Lipofectamine RNAi max per well diluted in optiMEM and siRNAs were diluted in optiMEM to reach the working concentration indicated in supplemental table 1. The following procedure and incubation times were as for Lipofectamine 2000. Besides the siRNA against the cellular proteins of interest, the AllStars negative siRNA (ctrl.) was included as a non-targeting control, whereas the AllStars death siRNA was used to test for successful transfection. Moreover, an siRNA targeting GFP was included to suppress the expression of the HPV16-GFP reporter plasmid as a measure for maximal reduction of infection. For RNAi against WASH, cells were transfected twice in 48 h intervals. Cells were infected with HPV16-GFP at 48 h post transfection to result in 20% infection in ctrl. negative transfected controls. In 12-well plates, infection was performed and analyzed by flow cytometry as described above. Absolute infection values were normalized to ctrl. transfected controls. In 96-well plates, the virus was added without prior medium exchange to reduce cell loss. At 48 h p.i., cells were fixed in 4% PFA in PBS and nuclei were stained with RedDot2 for 30 min after permeabilization with 0.1% Triton in PBS. Infection was analyzed by automated microscopy on a Zeiss Axiovert Z.1 microscope equipped with a Yokogawa CSU22 spinning disk module (Visitron Systems). Images were acquired using a 20x objective, a CoolSnap HQ camera (Photometrics) and MetaMorph Software. Cell number and infection were determined using a MATLAB-based infection scoring procedure (Engel et al., 2011). The program detects nuclei and infection signal individually, based on their limiting intensity edges. The edges were filled to objects, which were classified by size. Binary masks of nuclei and infection signal were created and cells were classified as infected if equal or greater than 5% of their nuclei overlapped with infection signal above a certain threshold. In this study, signal twice above the background in the uninfected sample was considered infected. An infection index was obtained for each image and averaged per well (Snijder *et al*., 2012).

Infection with VV-GFP was carried out in 96-well plates following the same protocol as for HPV16 with the exception that 3 x 10^3^ cells were transfected. Virus amounts leading to 20% infection in ctrl. treated controls were used. Cells were fixed at 6 h p.i. and analyzed by automated microscopy, as described above.

For siRNA experiments with SFV, 5 x 10^4^ HeLa Kyoto cells were reverse transfected in 12-well plates. Infection was performed by addition of the virus to infection medium (RPMI supplemented with 10% FBS, 10 mM HEPES (pH 7.3)). The virus was bound on a shaker at 37°C. At 2 h p.i., the inoculum was replaced by growth medium. Cells were trypsinized and fixed at 6 h p.i.. Since SFV did not carry a fluorescent reporter plasmid, samples were immunostained for SFV E1/E2 after fixation at 6 h p.i.. Cells were permeabilized with FACS perm (250 mM EDTA, 2% FBS, 0.02% NaN_3_, 0.05% Saponin in PBS) for 30 min at room temperature and subsequently incubated with the Lady Di antiserum (Singh and Helenius, 1992) diluted in FACS perm for 2 h at room temperature. Samples were washed thrice with FACS perm and incubated with an anti-rabbit AF488 secondary antibody in FACS perm for 1 h at room temperature. Washing with FACS perm was followed by infection scoring with FACS analysis (FACSCalibur, Becton Dickinson) as described for HPV16. Infection values were normalized to crtl. using Microsoft Excel.

##### Infection studies in transiently transfected cells

NIH-3T3 wild type and WASH KO cells were seeded in 12-well plates (4 x 10^4^ cells/well). One day later, cells were transfected with plasmids (1 µg) encoding EGFP-WASH or EGFP, YFP-WASH, YFP-WASH dWCA, YFP-WASH K220R, YFP-WASH Y141F or YPF-WASH Y141E using Lipofectamine 2000 (0.5 µl/well) diluted in optiMEM. Incubation times were the same as for virus preparation. At about 16 h post transfection, cells were infected with HPV16-RFP as described above to result in 20% infection in control cells transfected with GFP. Cells were trypsinized at 48 h p.i., fixed in 4% PFA in PBS and analyzed by flow cytometry (Guava easyCyte, Merck). Final analysis was performed with FlowJo. Transfected cells were gated with help of untransfected controls. Then, the GFP positive population was gated for infection using transfected, but uninfected controls. The percentage of transfected and infected cells (GFP and RFP positive) was normalized to NIH-3T3 wild type cells transfected with the GFP control to obtain relative infection values using Microsoft Excel.

HeLa ATCC cells were transfected with Scar-W and -WA constructs as described above 16-24 h prior to infection. Cells were infected with HPV16-RFP and fixed 48 h p.i. using 4% PFA. Infection was scored using an Olympus IX70 inverted microscope equipped with an electron multiplier CCD camera (EDMCCD, C9100-02, Hamamatsu Photonics K. K.) and a monochromator for epifluorescence excitation. Images were thresholded manually and at least 100 cells were scored for transfection and infection using Fiji.

##### HPV16 binding assay

For analysis of HPV16 binding by flow cytometry, 5 x 10^4^ NIH-3T3 wild type and WASH KO cells were seeded per well of a 12-well plate. The next day, fluorescently labeled HPV16-AF488 (∼1000 particles/cell) was bound to the cells for 2 h on a shaker at 37°C. Cells treated with siRNAs were reseeded at 48 h post transfection (5 x 10^4^ cells/well). Virus binding was performed once cells were attached, typically about 6 h post seeding. As a non-binding control, HPV16-AF488 was pre-incubated with 1 mg/ml heparin for 1 h at room temperature prior to binding to cells (Cerqueira *et al*., 2013). Cells were trypsinized and fixed with 4% PFA. Virus binding was analyzed by measuring the mean fluorescence intensity (geometric mean) of cells in flow cytometry (FACSCalibur). The geometric mean of uninfected cells was subtracted from infected cells and virus binding was normalized to control cells. A similar procedure was applied to measure virus binding to HeLa Kyoto cells depleted of SNX2 or SNX5. Binding was qualitatively assessed for NIH-3T3 wild type and WASH KO cells. For this, HPV16-AF568 was bound as described above. At 2 h p.i., cells were fixed with 4% PFA and stained with 0.1 µg/ml phalloidin-Atto488 diluted in PHEM buffer (60 mM PIPES, 10 mM EGTA, 2 mM MgCl_2_, 25 mM HEPES, pH 6.9) supplemented with 0.01% Triton X-100 for 30 min. Cells were washed thrice with PBS and mounted on glass slides using AF1 mounting medium. Images were acquired with a Zeiss Axiovert Z.1 microscope equipped with a Yokogawa CSU22 spinning disk module (Visitron Systems) using a 40x objective, a CoolSnap HQ camera (Photometrics) and MetaMorph Software. Z-stacks covering the cell volume were converted to maximum intensity projections using Fiji. Brightness and contrast were adjusted using uninfected samples.

##### Infectious internalization assay

One day prior to infection, 5 x 10^4^ HeLa Kyoto cells were seeded per well of a 12-well plate. Cells were pre-incubated with inhibitors and infected with HPV16-GFP as described above. At 12 h p.i., extracellular virus was inactivated by washing with 1 ml 0.1 mM CAPS buffer (pH 10.5) for 90 sec (Schelhaas *et al*., 2012; Becker *et al*., 2018). The cells were washed twice with PBS to remove CAPS and infection was continued in fresh growth medium. To control for inhibitor reversibility, the inhibitor was washed out thrice with PBS without prior CAPS treatment and fresh growth medium was added. At 48 h p.i., cells were fixed and infection was scored by flow cytometry as described above. Infection results were normalized to inhibitor reversibility.

##### Endocytosis assay with HPV16-pHrodo

Cells were reverse transfected with siRNAs as described above. At 48 h after the first (Arp3, SNX2) or second (WASH) siRNA transfection, HPV16-pHrodo (∼1000 particles/cell) was added to 350 µl growth medium and bound for 2 h on a shaker. The inoculum was replaced at 2 h p.i. and cells were imaged live at 6 h p.i. at 37°C and 5% CO_2_ in humidified atmosphere using custom made imaging chambers and DMEM high glucose without phenol red supplemented with 10% FBS, 1% L-glutamine and 1% penicillin/streptomycin. Images were acquired on a Zeiss Axiovert Z.1 microscope equipped with a Yokogawa CSU22 spinning disk module (Visitron Systems) using a 40x objective, a CoolSnap HQ camera (Photometrics) and MetaMorph Software. Average intensity projections of confocal slices were generated using Fiji software. Intensity based analysis was performed with CellProfiler (Becker *et al*., 2018). In brief, the virus signal was enhanced by application of a white top-hat filter. Virus spots were segmented by application of a gaussian filter and maximum correlation thresholding. Virus intensity was then measured in enhanced and original images. Pivot tables (Microsoft Excel) were used to summarize the intensity of spots per condition. These values were normalized to the cell number, which was determined by manual counting from brightfield images. The virus intensity per cell was then normalized to ctrl. treated controls to obtain relative internalization values. Cell outlines were created manually for presentation purposes.

#### Electron microscopy and CLEM

##### CLEM

A circular mark used for localization of unroofed cells was generated in the center of coverslips (22 mm diameter) using a diamond knife. For ECM production, 2 x 10^6^ HaCaT cells were seeded onto the coverslips placed in a 6-well plate. At 48 h post seeding, ECM was obtained by detaching cells through incubation with 10 mM EDTA/EGTA for 45 min at 37°C, subsequent clapping of the plate and washes with PBS (Culp et al., 2006). HPV16-AF568 was bound to the ECM in 1 ml growth medium/well for 2 h on a shaker at 37°C. HaCaT cells were trypsinized and 12 x 10^5^ cells/well were seeded onto the virus bound to ECM. At 1 h post seeding, 10 µg/ml cytoD or DMSO (solvent control) were added. A total of 6 h post seeding, cells were put on ice and washed thrice with cold stabilization buffer (70 mM KCl, 30 mM HEPES (pH 7.4 with KOH), 5 mM MgCl_2_). For unroofing, the cells were kept on ice and 1 ml cold 2% PFA in stabilization buffer was aspirated with a 1 ml pipette. The pipette was positioned above the marked area in the center of the coverslip and PFA was harshly released onto the cells. The coverslip was then rapidly transferred to a new well containing cold 2% PFA in stabilization buffer to avoid sedimentation of cell debris on the unroofed membrane. Membrane sheets were fixed for 10 min at 4°C. Samples were mounted in custom-made imaging chambers and imaged in PBS at a Nikon Ti Eclipse microscope equipped with a PerkinElmer UltraVIEW VoX spinning disk module. Images were acquired using a 60x objective, an Orca Flash 4 camera (Hamamatsu) and Volocity software (PerkinElmer, version 6.3). Montages of 10 x 10 images and 10% overlap were acquired around the center of the marked area. The unroofed membranes were prepared for EM by fixation with 2% glutaraldehyde (GA) in PBS overnight at 4°C. After two washes with water, samples were incubated with 0.1% tannic acid for 20 min at room temperature and subsequently washed with water. Contrasting was performed with 0.1% uranyl acetate (UAC) for 20 min at room temperature. After three washes with water, samples were dehydrated with a series of ethanol solutions (15%/30%/50%/70%/80%/90%/100%). Coverslips were incubated with each solution for 5 min, incubation with 100% ethanol was repeated thrice. Samples were dried using hexamethyldisilazane (HDMS). After 5 min incubation at room temperature, fresh HDMS was added and samples were incubated for further 30 min at room temperature. Coverslips were dried and coated under vacuum using a Balzers BAF301 device (former Balzers AG, Liechtenstein). A first layer of platinum was applied at an angle of 11°C while rotating. A second layer of carbon was applied at an angle of 90° while rotating. Coverslips were cut to fit on EM grids before 5% hydrofluoric acid were used to separate the metal replica from the glass. Replicas were extensively washed with water prior to transfer to glow discharged, formvar coated EM grids. Replicas were imaged with a phase contrast microscope for orientation. Intact membranes associated with virus particles were manually selected based on overlays of images from fluorescence and phase contrast microscopy and imaged at a transmission EM (Jeol JEM-1400, Jeol Ltd., Tokyo, Japan, camera: TemCam F416; TVIPS, Gauting, Germany). Membrane sheets were imaged using montages of 5 x 5 images and 15% overlap. Fluorescence and EM images were initially overlaid manually using Photoshop, then the Fiji plugin Landmark Correspondences (Saalfeld and Tomancák, 2008) was used for transformation of the fluorescence image according to the EM image using three manually identified landmarks. For analysis, HPV16 was identified manually based on the fluorescent signal and classification was done by visual evaluation of associated structures in transmission EM images. At total of 134 and 101 membrane associated virus particles in DMSO (7 membranes) and cytoD (5 membranes) treated cells were analyzed, respectively.

##### Ultra-thin section EM

Samples for ultra-thin section EM were prepared and analyzed as described previously (Bannach *et al*., 2020). A total of 1-2 x 10^5^ NIH 3T3 wild type, WASH KO and HeLa Kyoto cells or were seeded in 3 cm dishes. Two days post seeding, cells were either pretreated with inhibitors for 30 min or were left untreated. Cells were infected with 40 µg HPV16 PsVs in 1 ml growth medium. At 6 h p.i., cells were fixed in 2.5% GA in PBS (pH 7.2) for 10 min at room temperature. A second fixation was performed with cold 2.5% GA at 4°C overnight. Cells were washed thrice with PBS, post-fixed with 1% OsO_4_ in ddH_2_O for 1 h and washed twice with ddH_2_O at room temperature and at 4°C for 20 min, respectively. Block contrasting was performed with 0.5% UAC in ddH_2_O at 4°C overnight. Cells were washed thrice with ddH_2_O and dehydrated using ascending graded alcohol series. Detaching and dehydrating with propylene oxide were followed by incubation with propylene oxide and epoxy resin (1:3, 1:1, 3:1) for 2 h each, before cells were incubated with pure epoxy resin for 3 h and embedded in BEEM capsules. The resin was allowed to polymerize at 60°C for three days before 60 nm ultra-thin sections were cut and counterstained with uranyl acetate and lead. Samples were imaged at a 80 kV on a Tecnai 12 electron microscope (FEI) using an Olympus Veleta 4k CCD camera or Ditabis imaging plates. Images were contrast enhanced and cropped using Adobe Photoshop CS4. The total number of endocytic pits per cell was determined for 31 and 43 cells in wild type and WASH KO cells, respectively, in two independent experiments. Only endocytic pits containing virus(es) were counted, since HPV16 pits are hardly distinguishable from uncoated pits from other endocytic pathways without further staining. Pit numbers were normalized to wild type cells.

##### Immunogold labeling

HeLa ATCC cells (2-3 x 10^5^ cells) were seeded in 6 cm dishes. The next day, cells were transfected with a plasmid encoding EGFP-WASH or EGFP-SNX2 (7 µg) using Lipofectamine 2000 (3.5 µl) according to the manufacturer’s instructions or left untransfected. At 48 h post transfection, cells were infected with 80 µg HPV16 PsVs and incubated for 6 h at 37°C until pre-fixation by addition of 4% formaldehyde in 0.1 M phosphate buffer (pH 7.2) to the culture medium (1:1 ratio) for 5 min. Then, cells were fixed in 2% formaldehyde, and 0.2% GA in 0.1M phosphate buffer (pH 7.2). Samples were processed for transmission EM as previously described (Humbel and Stierhof, 2009). In brief, cells were quenched by incubation in 0.1% glycine in 0.1 M PB (2 x 30 min), washed thrice in 0.1 M PB for 30 min and scraped with 1% gelatin. After centrifugation, the gelatin was replaced with 12% gelatin and cells were infused at 37°C. Cells were cooled down on ice and the gelatin-cell pellet was cut into small blocks that were infused with 2.3 M sucrose overnight at 4°C. The blocks were mounted on specimen carriers and frozen in liquid nitrogen. Ultra-thin cryosections were prepared according to Tokuyasu (Tokuyasu, 1980). In brief, ultra-thin cryosections of 50 nm thickness were prepared with an EM UC6/FC6 ultramicrotome (Leica Microsystems). Sections were collected in a sucrose-methylcellulose mixture and stored on transmission EM grids at 4°C until further processing. Methylcellulose was melted and sections were washed five times with 20 mM glycine in PBS. Quenching was followed by blocking with 1% BSA for 3 min. Cells were then incubated with a GFP-antibody for 30 min, washed 6 times with 0.1% BSA in PBS and incubated with protein A gold 15 nm for 20 min. Sections were rinsed 10 times with PBS and re-fixed in 1% GA in PBS (pH 7.2). For double labeling, the sections were quenched, blocked and immunostained with an actin antibody as described above. Then sections were incubated with a rabbit anti-mouse bridging antibody followed by 6 washes with 0.1% BSA in PBS and incubated with protein A gold 10 nm (1:50). Sections were rinsed 10 times with PBS and re-fixed in 1% GA in PBS, pH 7.2. Sections were rinsed 10 times with ddH_2_O and contrasted with uranyl acetate for 6 min (pH 7). After one wash with ddH_2_O, cells were embedded in an uranyl acetate-methylcellulose mixture (pH 4) for 10 min. After looping out with filter paper, sections were dried and images were acquired as above.

For quantification, 50 cell profiles were randomly selected. Antibody signals were determined as the number of gold particles associated with specific cellular compartments. Gold particles counted in 50 profiles of HeLa cells treated with Lipofectamine 2000 (negative control) were used as control.

#### Fluorescence microscopy

##### CLC and virus internalization analysis by live cell TIRF microscopy

HeLa ATCC cells were seeded on coverslips in 12-well plates (5 x 10^4^ cells/well) one day prior to transfection. Cells were transfected with plasmids encoding lifeact-EGFP, EGFP-WASH, EGFP-SNX2, mRFP-CLC or EGFP-Dyn2 as described above. For internalization analysis, fluorescently labeled virus (HPV16-AF594/AF647) was bound at 37°C at about 18 h post transfection. At 1 h p.i., coverslips were mounted in custom-made imaging chambers. Cells were imaged at 37°C and 5% CO_2_ in humidified atmosphere in DMEM without phenol red supplemented with 10% FBS, 1% L-glutamine and 1% penicillin/streptomycin. Time lapse movies of cells expressing lifeact-EGFP were acquired with a 60x TIRF-objective on an Olympus IX70 microscope equipped with a TIRF condenser and an electron multiplier CCD camera (EDMCCD, C9100-02, Hamamatsu Photonics K. K.) using MetaMorph software (Molecular Devices) (Visitron Systems). All other time lapse movies were acquired using a 100x TIRF-objective at an Olympus IX83 microscope equipped with a four-line TIRF condenser and an EMCCD camera (iXon Ultra 888, Andor Oxford Instruments) using CellSens Dimensions software (Olympus). Movies were acquired with 0.5 Hz frame rate for 5 min. HPV16 entry events were identified manually and the intensity of fluorescent proteins at virus entry sites was analyzed after rolling ball background subtraction and filtering (mean intensity filter) with Fiji. Kymographs, intensity profiles along a manually drawn line through the virus/clathrin signal over time, were generated with Fiji after background subtraction and filtering. Intensity profiles measured with Fiji using a circular region of interest, were processed by min/max normalization and aligned by setting the half time of virus internalization to 0 s. Profiles were plotted with Microsoft Excel. Moving averages of signals are shown as a trendline (period 4-20). The time points of recruitment onset, maximal signal or signal loss were manually determined relative to the half time of internalization and box plots were generated with GraphPad Prism. Cells co-transfected with CLC and Dyn2 were analyzed the same way at about 16 h post transfection. Movies were compressed to 20 fps and PNG.

##### Proximity ligation assay

For ECM production on coverslips, 5 x 10^5^ HaCaT cells were seeded per well of a 12-well plate and cultivated at 37°C for 2 days. Additionally, 7 x 10^4^ HaCaT cells were seeded per well and transfected the next day with a plasmid encoding HA-CD151. The procedure was the same as for infection studies with transfected cells, but 0.4 µg DNA and 1 µl Lipofectamine 2000 per well were used. ECM on coverslips was obtained by detaching cells with 0.5 ml 10 mM EDTA/EGTA as described above. HPV16-AF488 (∼1000 particles/cell) was bound to the ECM in 400 µl growth medium on a shaker at 37°C. At 2 h post binding, HaCaT cells expressing HA-CD151 were trypsinized and transferred onto the virus bound to ECM. Cells were allowed to attach for 5 h and fixed in 2% PFA in PBS for 10 min at 4°C. Cells were permeabilized with 0.2% Brij 58 in PBS for 10 min at room temperature prior to blocking in 1% BSA in PBS for 30 min at room temperature. Primary antibody staining against HA (1:10,000) and WASH (1:500) was carried out in a wet chamber overnight at 4°C. Next, cells were incubated with anti-mouse PLUS and anti-rabbit MINUS Duolink PLA probes diluted 1:5 in 1% BSA in PBS for 1 h at 37°C in a humidity chamber. Duolink In Situ Detection Reagents Red were used for further sample processing. For ligation of PLA probes, 1.25 U ligase per sample were added to the corresponding ligation buffer diluted in ddH_2_O. The cells were incubated in at wet chamber for 30 min at 37°C followed by two washes with wash buffer B (2 min, room temperature). Amplification was performed with 6.25 U polymerase and Duolink Amplification Red (1:5) diluted in ddH_2_O in a wet chamber at 37°C for 100 min. Cells were washed twice for 10 min with wash buffer B at room temperature followed by a quick wash with 0.1x wash buffer B in ddH_2_O. As a counterstain, HA (CD151) was detected with an anti-mouse AF647 antibody, which was diluted 1:2000 in 1% BSA in PBS. Samples were transferred to custom made imaging chambers. Images were acquired with a 100x TIRF-objective at an Olympus IX83 microscope equipped with a four-line TIRF condenser and an EMCCD camera (iXon Ultra 888, Andor Oxford Instruments) using CellSens Dimension software (Olympus).

#### Flow cytometry analysis of receptor cell surface levels in U2OS WASH KO cells

U2OS WT and WASH KO cells were detached by incubation with 3 ml 50 mM EDTA for 10 min at 37°C. After fixation with 2% PFA (4°C, 10 min), cells were sedimented (5 min, 600 x g, 4°C) and washed twice with PBS. Blocking was performed with 1% BSA for 1 h at room temperature. Subsequently, 7 x 10^4^ cells were incubated with anti EGFR (1:150) or CD151 (1:500) antibodies diluted in 1% BSA at 4°C overnight. Three washes with PBS were followed by incubation with Alexa Fluor 488 conjugated secondary antibodies diluted in 1% BSA for 1 h at room temperature. Cells were washed thrice with PBS and resuspended in 300 µl flow cytometry buffer (250 mM EDTA, 2% FBS, 0.02% NaN3 in PBS). A Gallios Flow Cytometer (Beckman Coulter) was used to determine receptor cell surface levels as the geometric mean fluorescence intensity. Background values from controls incubated with secondary antibodies only were subtracted and intensities were normalized to wild type cells.

#### Quantification and statistical analysis

Information on data representation (mean ± SEM) and n can be found in the figure legends. Statistical significance was determined using unpaired t-tests conducted with GraphPad Prism. A P-value below 0.05 (P < 0.05) indicated a significant difference, denoted by asterisks in the figures (* P ≤ 0.05, ** P ≤ 0.01, *** P ≤ 0.001). If not specifically indicated, differences were not significant.

---

**Supplemental table 1.** (siRNA)

| siRNA | working concentration (nM) | Qiagen identifier |
| --- | --- | --- |
| AllStars death | 10 | SI04381048 |
| AllStars negative (siCtrl.) | 10/20/50 | SI03650318 |
| Arp3 siRNA_1 | 10 | Hs_ACTR3_5 |
| Arp3 siRNA_2 | 10 | Hs_ACTR3_8 |
| CCDC53 siRNA_1 | 10 | Hs_CCDC53_3 |
| CCDC53 siRNA_2 | 10 | Hs_CCDC53_4 |
| CtBP1 siRNA_1 | 10 | Hs_CTBP1_5 |
| CtBP1 siRNA_2 | 10 | Hs_CTBP1_6 |
| DIAPH2 siRNA_1 | 10 | Hs_DIAPH2_1 |
| DIAPH2 siRNA_2 | 10 | Hs_DIAPH2_4 |
| DIAPH2 siRNA_3 | 10 | Hs_DIAPH2_6 |
| FAM21 siRNA_1 | 10 | Hs_FAM21_11 |
| FAM21 siRNA_2 | 10 | Hs_FAM21_12 |
| FNL2 siRNA_1 | 10 | Hs_FMNL2_6 |
| FNL2 siRNA_2 | 10 | Hs_FMNL2_7 |
| FNL2 siRNA_3 | 10 | Hs_FMNL2_8 |
| FNL3 siRNA_1 | 10 | Hs_FMNL3_1 |
| FNL3 siRNA_2 | 10 | Hs_FMNL3_5 |
| FNL3 siRNA_3 | 10 | Hs_FMNL3_6 |
| Formin 1 siRNA_1 | 10 | Hs_FMN1_5 |
| Formin 1 siRNA_2 | 10 | Hs_FMN1_6 |
| Formin 1 siRNA_3 | 10 | Hs_FMN1_7 |
| Formin 2 siRNA_1 | 10 | Hs_FMN2_12 |
| Formin 2 siRNA_2 | 10 | Hs_FMN2_7 |
| Formin 2 siRNA_3 | 10 | Hs_FMN2_9 |
| GFP siRNA | 10 | SI04380467 |
| JMY siRNA_1 | 50 | Hs_JMY_1 |
| JMY siRNA_2 | 50 | Hs_JMY_5 |
| N-WASP siRNA_1 | 10 | Hs_WASL_1 |
| N-WASP siRNA_2 | 10 | Hs_WASL_6 |
| RME8 siRNA_1 | 50 | Hs_DNAJC13_1 |
| RME8 siRNA_2 | 50 | Hs_DNAJC13_4 |
| RME8 siRNA_3 | 50 | Hs_DNAJC13_5 |
| RME8 siRNA_4 | 50 | Hs_DNAJC13_6 |
| strumpellin siRNA_1 | 10 | Hs_KIAA0196_7 |
| strumpellin siRNA_2 | 10 | Hs_KIAA0196_8 |
| SWIP siRNA_1 | 10 | Hs_KIAA1033_5 |
| SWIP siRNA_2 | 10 | Hs_KIAA1033_7 |
| WASH siRNA_1 | 20 | Hs_WASH1_4 |
| WASH siRNA_2 | 20 | Hs_WASH1_8 |
| WAVE1 siRNA_1 | 20 | Hs_WASF1_3 |
| WAVE1 siRNA_2 | 50 | Hs_WASF1_4 |
| WAVE2 siRNA_1 | 10 | Hs_WASF2_5 |
| WAVE2 siRNA_2 | 10 | Hs_WASF2_6 |
| WHAMM siRNA_1 | 50 | Hs_WHDC1_1 |
| WHAMM siRNA_2 | 50 | Hs_WHDC1_2 |
| Vps26 siRNA_1 | 10 | Hs_VPS26A_1 |
| Vps26 siRNA_2 | 10 | Hs_VPS26A_2 |
| Vps29 siRNA_1 | 10 | Hs_VPS29_6 |
| Vps29 siRNA_2 | 10 | Hs_VPS29_2 |

