## Supplementary figures and images for "Clathrin-independent endocytosis of Human Papillomaviruses is facilitated by actin nucleation promoting factor WASH"

### Figure S1

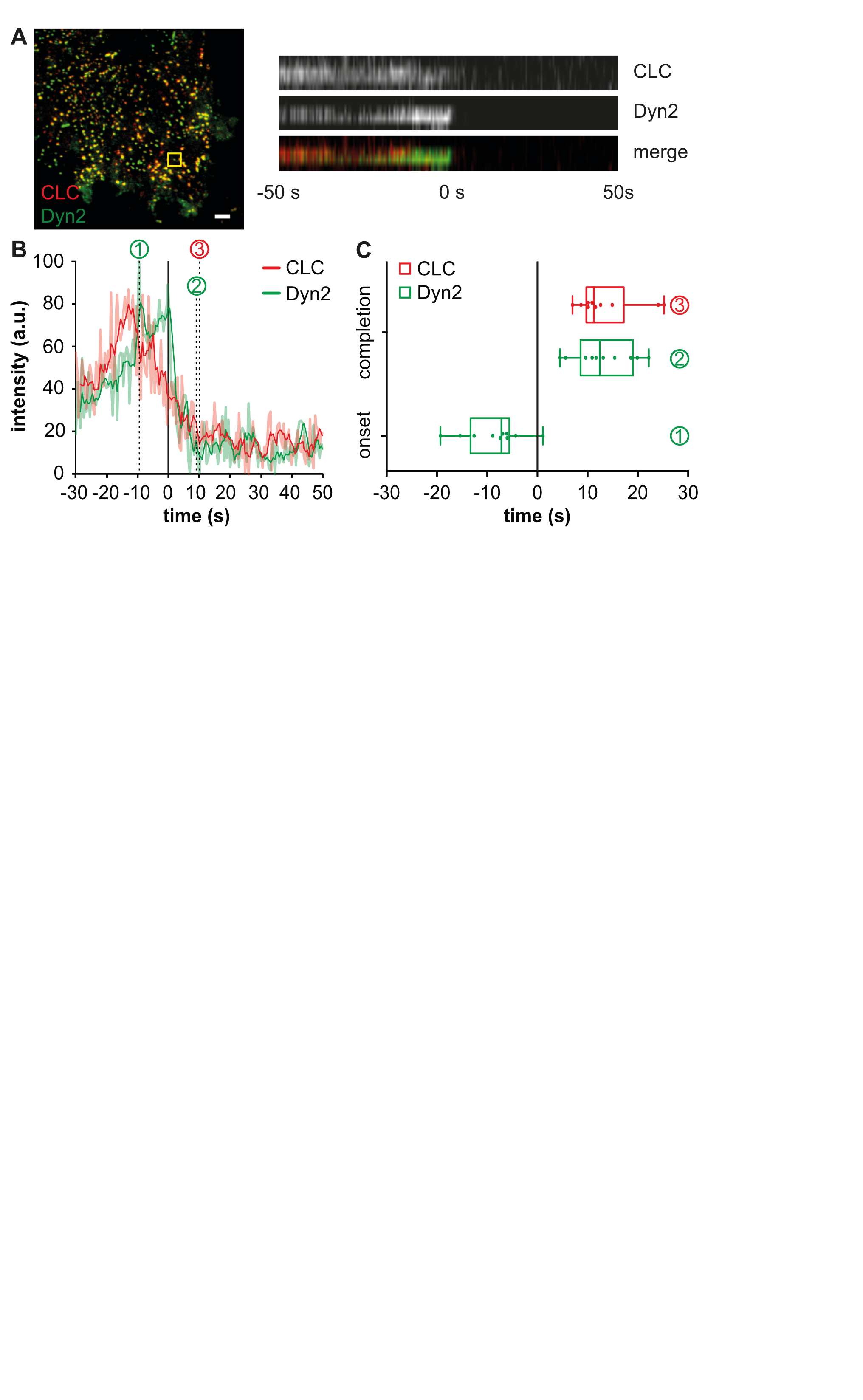

### Figure S2

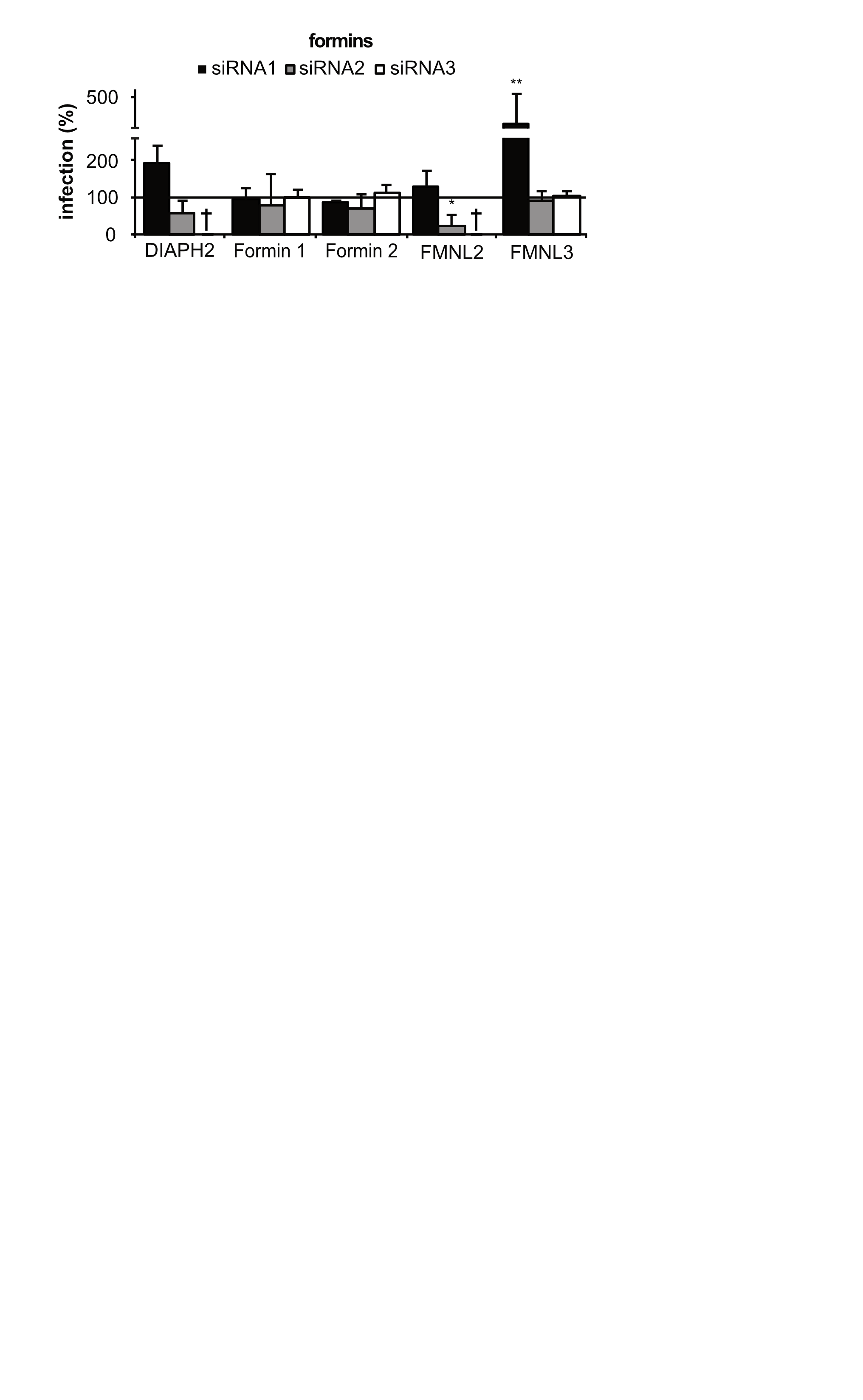

### Figure S3

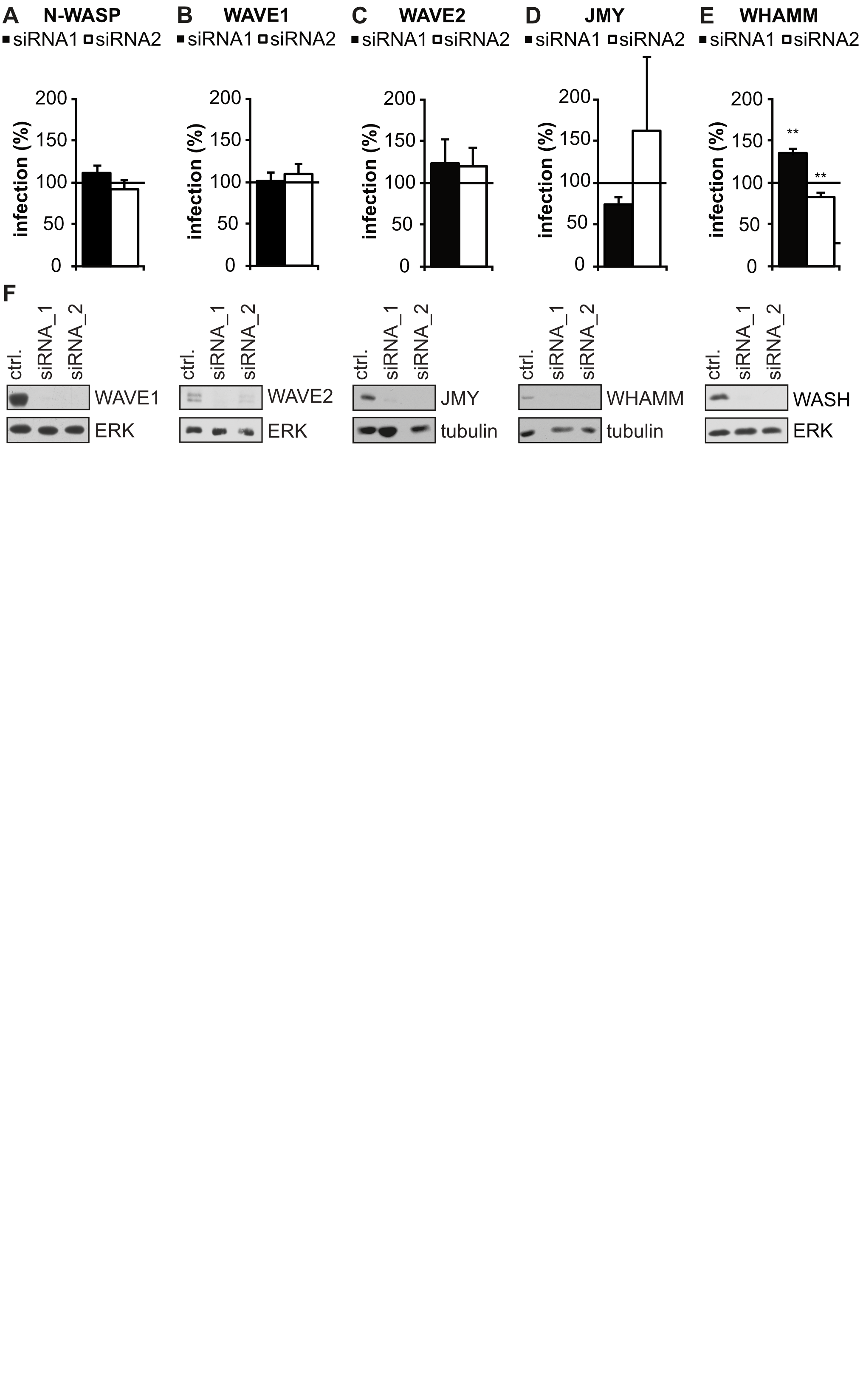

### Figure S4

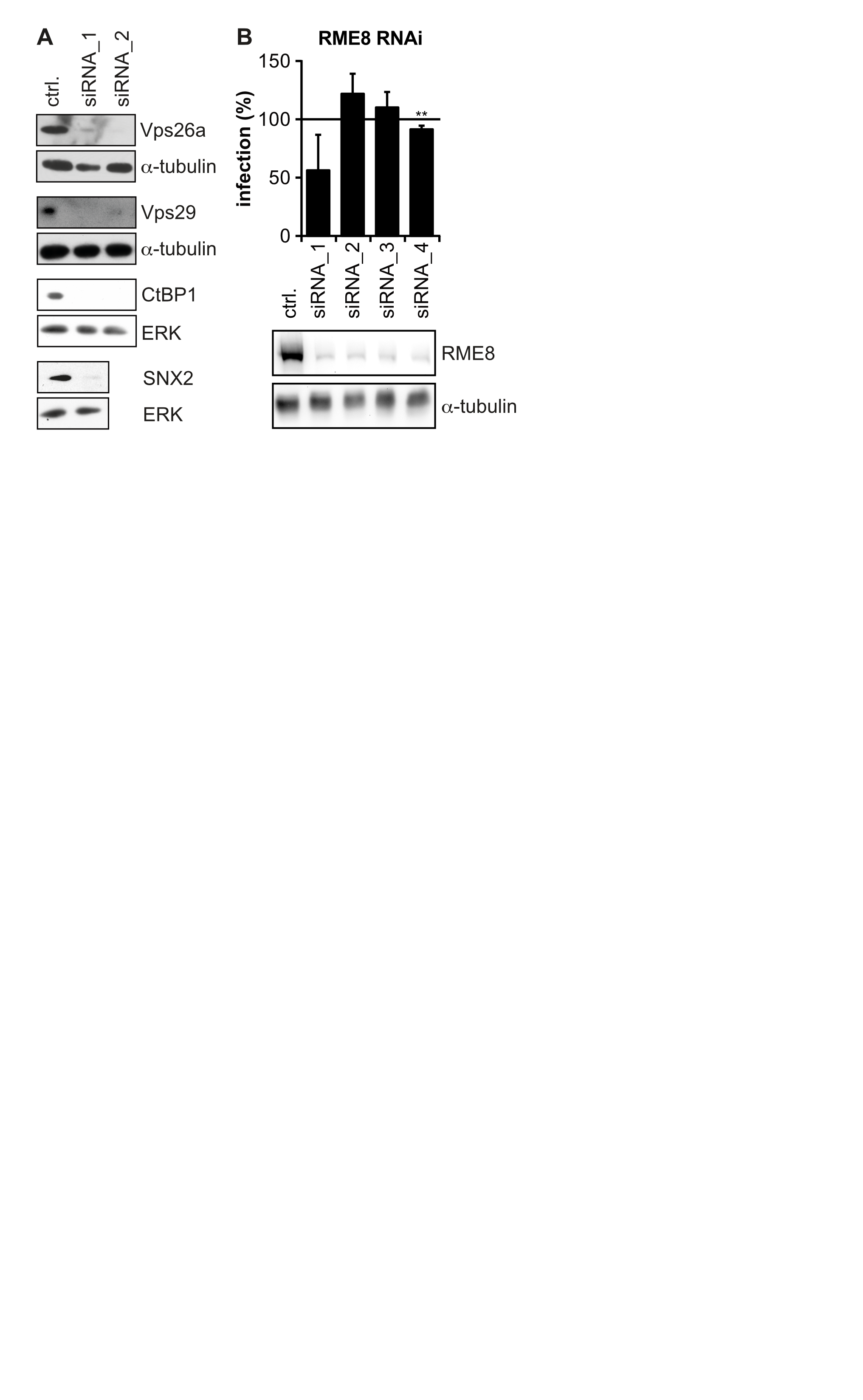

### Figure S5

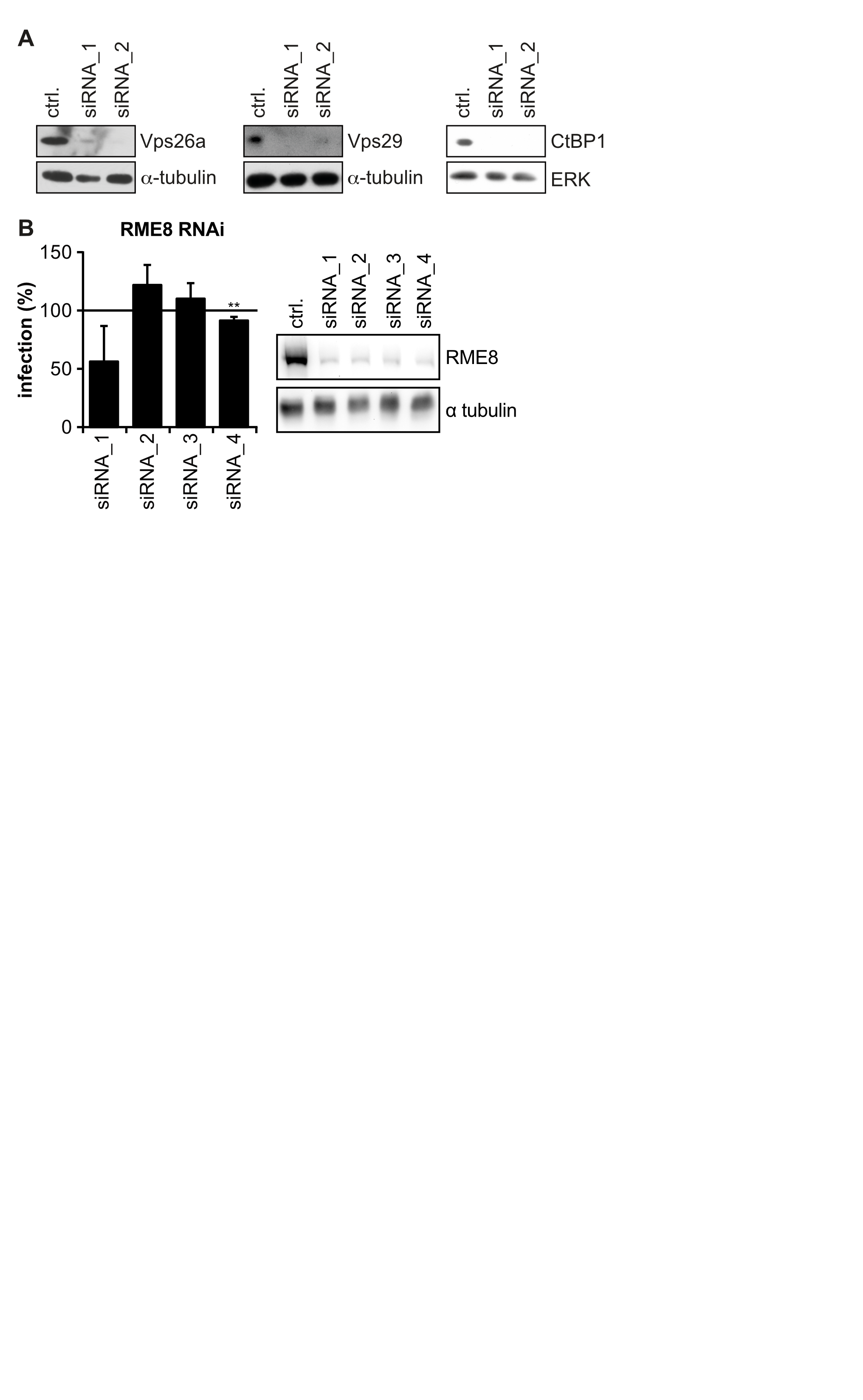

### Figure S6

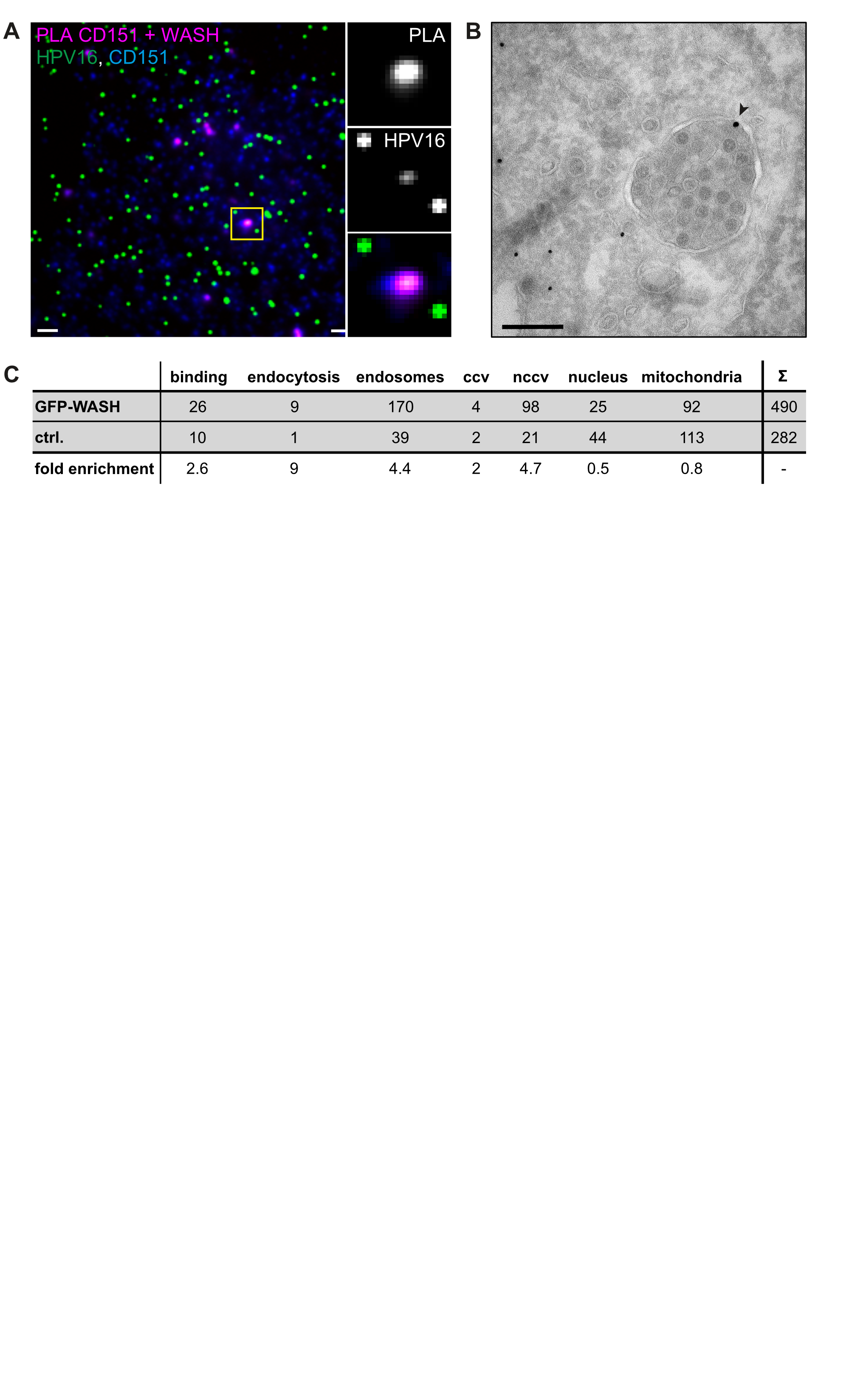
